# Ecological implications of removing a concrete gas platform in the North Sea

**DOI:** 10.1101/2020.04.16.044263

**Authors:** Joop W.P. Coolen, Oliver Bittner, Floor M.F. Driessen, Udo van Dongen, Midas S. Siahaya, Wim de Groot, Ninon Mavraki, Stefan G. Bolam, Babeth van der Weide

**Affiliations:** Wageningen Marine Research, P.O. Box 57, 1780 AB Den Helder, The Netherlands; Wageningen University, Aquatic Ecology and Water Quality Management Group, Droevendaalsesteeg 3a, 6708 PD Wageningen, The Netherlands; Bureau Waardenburg, P.O. Box 365, 4100 AJ Culemborg, The Netherlands; The Centre for the Environment, Fisheries and Aquaculture Science (Cefas), Lowestoft, NR33 0HT, UK

**Keywords:** Benthic biodiversity, epifouling, North Sea, artificial reef, gravity-based structure, gas platform

## Abstract

Artificial structures such as offshore oil and gas platforms can significantly alter local species communities. It has been argued that this effect should be considered during decisions over their removal during decommissioning. In the North Sea, leaving such structures in place is prohibited but derogations are allowed for large concrete installations. To assess removal options for one such installation, the Halfweg GBS (gravity-based structure) a concrete platform foundation off the Dutch coast, we studied the resident fouling macrofauna community. The faunal structure, biomass and trophic composition of the Halfweg was then compared with those from the surrounding seabed sediments, other local artificial structures and a natural rocky reef.

In total, 65 macrofaunal species were observed on the concrete (52 species), steel legs (32) and surrounding rock dump (44) of Halfweg. Mean Simpson diversity per sample was highest on the rock dump (0.71) but concrete (0.54) and steel (0.60) of the GBS were lower than seabed (0.69). Ten of the species observed on the concrete were not reported on other substrates while 10 of the species were also observed in the surrounding seabed. The GBS structure was numerically dominated by Arthropoda which comprised 98% of the total abundance. Mean ash free dry weight (AFDW) was significantly higher (p<0.001) on the Halfweg substrates (204 g AFDW per m^2^) than in the surrounding seabed (65 g AFDW per m^2^). Over 94% of the biomass on Halfweg consisted of the plumose anemone *Metridium senile*. While common on other reefs, this species was absent from the surrounding seabed. Macrofaunal feeding mechanisms of the concrete and rock dump communities on the GBS were similar to those of nearby sediments, although these differed from those on the Halfweg steel legs. Therefore, the presence of Halfweg alters the local community feeding modes. Multivariate analysis revealed that taxonomic structure of the GBS and other artificial structures significantly differed from that of the sedimentary habitats. Low numbers of non-indigenous species on Halfweg indicated that the structure does not act as a stepping stone for species invasions.

Our data show that the Halfweg structures significantly increase local biodiversity and biomass. Removal of the concrete and steel legs of the GBS (leaving the rock dump) will significantly reduce local macrofauna biodiversity. The long-term impact on macrofaunal biomass is low. Leaving the complete Halfweg structure in place will result in an enriched local macrofaunal biodiversity and feeding mode diversity.

## 1. INTRODUCTION

The presence of artificial structures such as oil and gas platform or wind turbine foundations in the marine environment induces significant changes on the local species diversity (Dannheim et al., 2020; Fowler et al., 2018). These *de facto* artificial reefs attract a community of epifouling species (Goddard and Love, 2010; Krone et al., 2013; Picken, 1986), fishes (Fujii, 2015; Love et al., 2003; Pradella et al., 2014) and mammals (Russell et al., 2014). Differences as well as similarities have been reported between artificial and natural reefs (Coolen et al., 2018; Dannheim et al., 2020; Wilhelmsson and Malm, 2008). In general, the presence of these installations is considered to increase local biodiversity (Dannheim et al., 2020) and population connectivity (Coolen et al., 2020; Henry et al., 2018; van der Molen et al., 2018). The structures have been suggested to act as stepping stones which can influence the distribution of native species (Friedlander et al., 2014) as well as non-native species (Yeo et al., 2010). Apart from their positive effects on biodiversity, artificial structures can have a positive impact on marine food webs. Studies have shown that fish species are attracted towards such structures due to the increased prey abundance, i.e. fouling fauna (Reubens et al., 2013, 2011). Furthermore, the introduction of scour protection layers increases the local food web complexity, supporting a high diversity of trophic levels (Mavraki et al., 2020a). The biodeposition processes of fouling organisms create organic-matter rich soft sediments near the base of offshore wind foundations, which in turn increases the abundance and species richness of the macrofaunal communities (Coates et al., 2014). Finally, fouling organisms are responsible for a negligible reduction of the local primary producers (Mavraki et al., 2020b). All these suggest that artificial structures could have beneficial effects on the local food web properties. The ecological importance of these structures should, therefore, be considered when decisions are taken regarding their removal as part of their decommissioning processes (Fowler et al., 2018).

Worldwide, a high number of oil and gas installations are scheduled to be decommissioned in the coming years (Fowler et al., 2018). During decommissioning, structures can be removed and brought ashore for scrapping or reuse (Schroeder and Love, 2004). In some cases, the structure foundations are left in place (Bull and Love, 2019) or relocated to be used as artificial reefs (Picken et al., 2000). In the North Sea region, the OSPAR decision on the Disposal of Disused Offshore Installations (OSPAR Commission, 1998), dictates that “*The dumping, and the leaving wholly or partly in place, of disused offshore installations within the maritime area is prohibited*”. However, derogations of this prohibition are allowed if assessment by the relevant competent authority “*shows that there are significant reasons why an alternative disposal [*..*] is preferable*”. One of the exclusion structures considered in this decision is the foundation of gravity based concrete installations.

In the Netherlands, the removal obligation for installations has been embedded in the Mining Act (Kingdom of the Netherlands, 2020) which states that an unused mining installation is to be removed. Currently, 160 oil and gas production installations are present within the Dutch North Sea (de Vrees, 2019). Most of these consist of steel jacket structures but a few Dutch installations are constructed using a concrete gravity-based foundation. To date, none of these concrete structures have been removed from the Dutch North Sea, and no derogations from the OSPAR decision have been proposed. In 2016, a proposal to leave two jacket foundations in place for 15 years to gain insights into the ecological effects of leaving structures in place after decommissioning (Maslin, 2016) was withdrawn for financial and permitting reasons. Thus, data regarding the ecological implications of removing such structures as opposed to leaving them in situ during decommissioning in this part of the North Sea is still lacking.

One of the few concrete foundations in Dutch waters is the Halfweg gravity-based structure (GBS). The Halfweg gas production platform was built in 1995 and operated by Petrogas E&P Netherlands B.V., the Netherlands (henceforth: Petrogas). The GBS was never used to store hydrocarbons and only functioned as a foundation to the platform. Gas production on the platform ended in 2016, followed by its decommissioning. Although the original intention was to remove the GBS together with its supporting steel legs and topside, a collision by a gas tanker vessel in December 2017 damaged the legs, preventing the lift of the whole structure at once (personal communication Alan Shand, Petrogas). Therefore, the topside and legs were removed from the GBS in January 2019 and the concrete structure remained on the seabed. Currently, Petrogas is evaluating options for the removal of the GBS and is considering four options. Three of these options involve the complete removal of the GBS using different methods to lift it, scattering approximately 50% of the surrounding rock dump in the area where the GBS is now, while the fourth option consists of leaving the whole GBS structure in situ.

To provide empirical data to aid decisions regarding the decommissioning of the GBS, we conducted a survey to acquire data to allow a comparison of the structural and functional (biomass and feeding modes) characteristics of the macrofaunal assemblages of the GBS structure and its associated rock dump with those of the surrounding sedimentary habitats. To place the ecological importance of the GBS into a wider context, these macrofaunal attributes were also compared with those of other artificial structures in the region including the steel piles of wind turbines and a natural rock reef. To aid this comparison the four removal options were reduced to two scenarios, these were:

1. The GBS is fully removed and the surrounding rock dump is partly scattered across the area, leaving the other part of the rock dump untouched;
2. The GBS is left in situ with the rock dump remaining unaltered.

## 2. MATERIALS AND METHODS

### 2.1 Study site

The Halfweg gravity-based structure (GBS) is located 26 km offshore from the Dutch coast (52.882°N 4.320°E, WGS84, Figure 1) in a water depth of 25 m.

**Figure 1:**
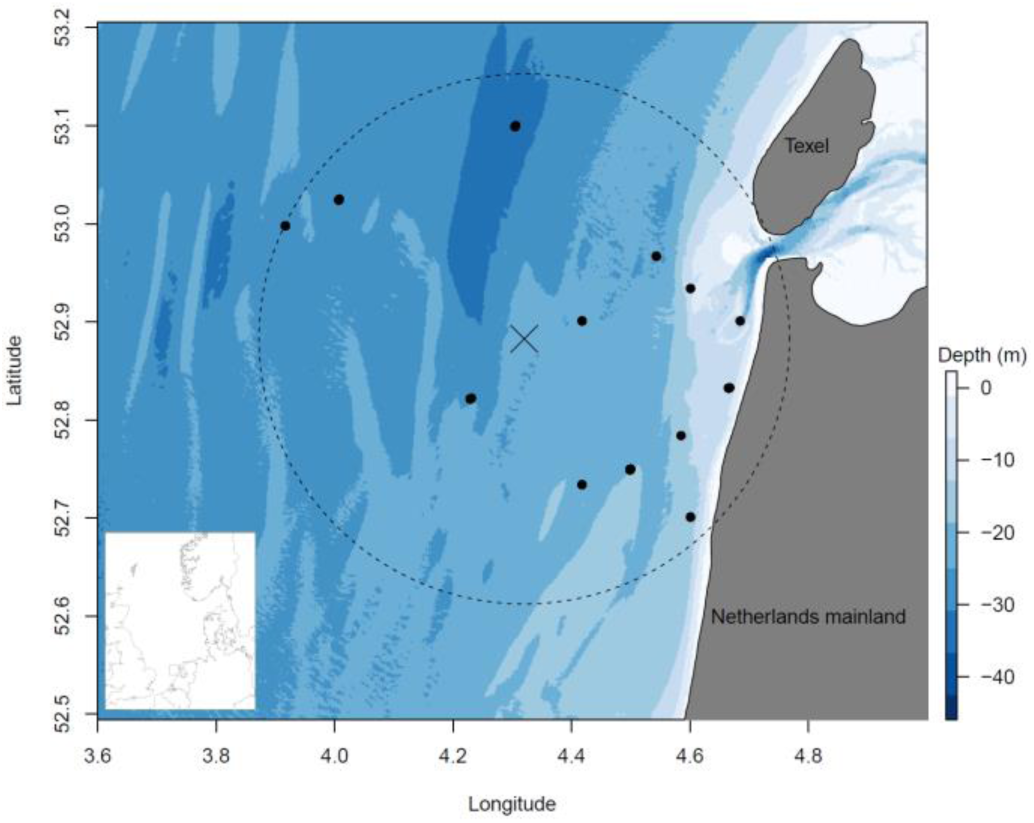
Sample locations. Map showing the Dutch coast, position of the Halfweg GBS structure (X) and locations of seabed data extracted from MWTL (black dots; Marine Information and Data Centre, 2019) within a 30 km radius from Halfweg (striped line). Seabed bathymetry (EMODnet, 2019) and depth in metres are provided in legend on the right. Inset map: Greater North Sea with rectangle indicating the study area. Latitude and longitude in decimal degrees (WGS84).

It is a concrete box-shaped structure of 26*28*6 m (width*length*height; Figure 2) providing a hard surface area available to macrofauna of approximately 1,376 m^2^. The walls of the GBS are almost vertical, with a slight outward slope, resulting in a base wider than the top.

**Figure 2:**
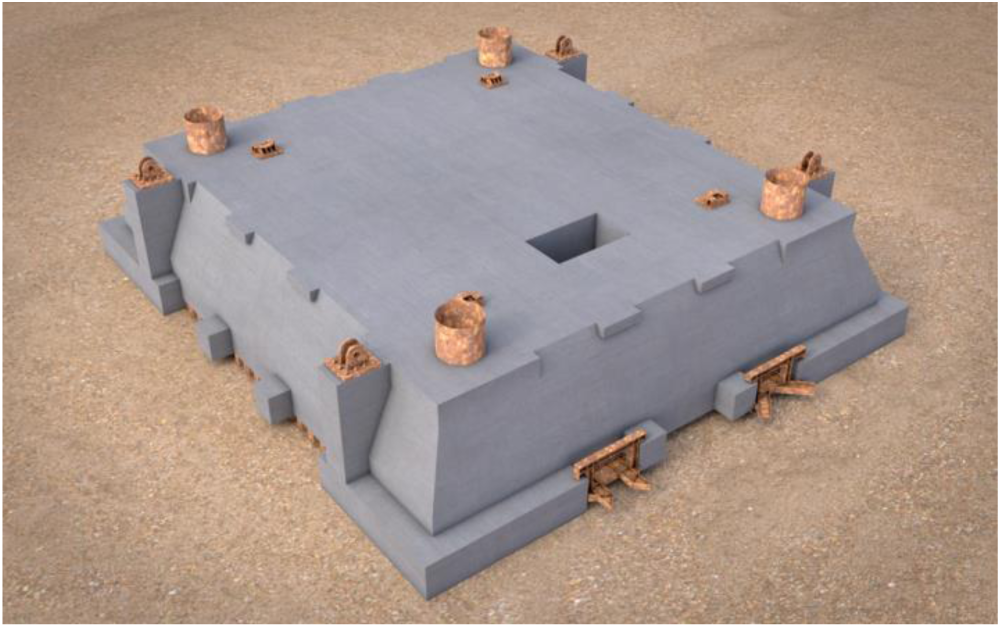
Halfweg gravity based foundation. Model representation of the concrete GBS as it currently lies on the seabed (rock dump not shown). Image provided by Petrogas E&P Netherlands.

It is placed on the seabed in a south-east to north-west orientation and it is surrounded by a rock dump of different sizes (10-70 cm diameter; Figure 3) in a radius of approximately 15-20 m from the GBS and a height of up to 6 m, covering a total seabed area of approximately 2,889 m^2^ (Figure S1). Since rock dumps form complex structures it was assumed that the hard substrate area available for macrofauna was larger than the area of seabed covered. The rock surface area available for macrofauna was calculated by generating 1,000,000 rocks of random size between 10-70 cm length, width and height, calculating the approximated surface area per rock (Graham et al., 1988), in comparison with the seabed covered by that rock, assuming a vertically projected rectangle shaped area (length*width of each rock). This ratio was averaged to 3.59 m^2^ rock per m^2^ seabed and multiplied by the 2,889 m^2^ of seabed covered. It was assumed that the top layer of rocks was fully available for macrofauna on all sides. Unavailability of the surface on rocks touching each other, additional surface available on rocks below the first layer as well as cover by sand of rocks on the edges of the rock dump area were not considered. The estimated total surface area of rocks available to macrofauna was 10,389 m^2^. Most hard surface present at Halfweg is composed of concrete and the rock dump, with a small steel surface available in the form of the remains of the 1.7 m diameter legs, which are between 2 and 3.5 m in height above the GBS (Figure 2). The hollow legs, which intrude 6 m into the GBS, are open on the top allowing their inner surface to provide a potential substratum for macrofaunal colonisation. Total steel surface area available for marine growth excluding the deeper, internal regions where limited water exchange results in oxygen- and nutrient-depleted waters, was estimated to be 120 m^2^.

**Figure 3:**
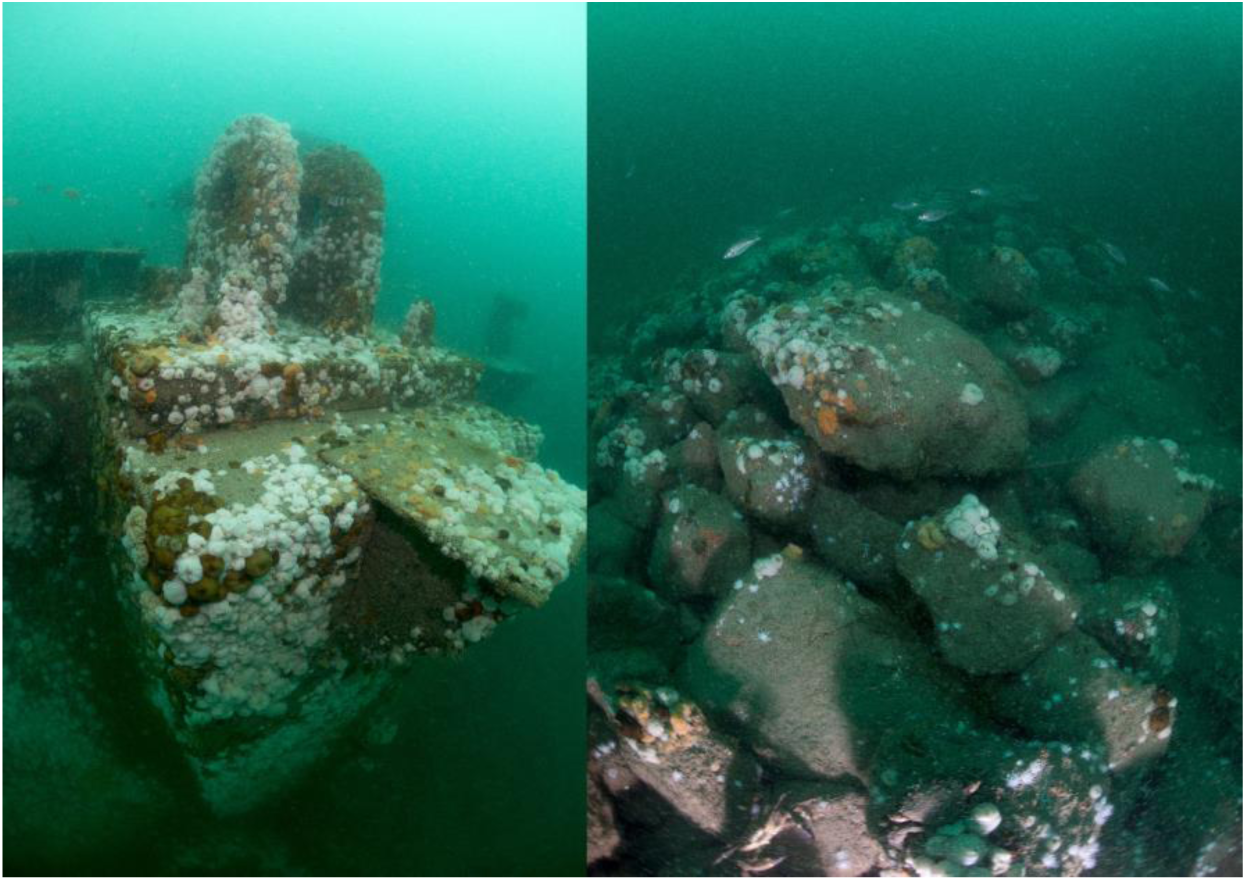
In situ impression of the Halfweg GBS and rock dump. Under water photographs of: left: a detail of the GBS showing one of the lifting points and surrounding concrete substrate; right: overview of the rock dump surrounding the GBS. Photos by Udo van Dongen (Bureau Waardenburg b.v.).

The spatial area of seabed occupied by the complete GBS structure is approximately 3,617 m^2^. The surrounding seabed habitats within a 30 km radius of the structure are largely composed of sand with patches of coarse substrate mostly located north of the GBS (EMODnet, 2020). Depth varies between 0 m to the east to 33 m in the west of the area (Figure 1). To the northeast the area borders the Wadden Sea, which is enclosed between islands (e.g. Texel; Figure 1) and the mainland. Several other gas platforms and shipwrecks are present in the region (Coolen et al., 2020 [Figure 1]).

### 2.2 Data acquisition

#### 2.2.1. Sampling of the Halfweg structure

In September 2019 surveys were conducted to acquire samples from the marine growth occurring on the GBS including the steel legs and the surrounding rock dump. A total of 39 samples were taken by commercial diving certified marine biologists using scuba equipment. Samples were collected along transect lines placed in four directions on the GBS (Figure 4). In each direction, two samples were taken from the horizontal top of the GBS, three samples from the vertical-diagonal side of the GBS and three from the rock dump. Finally, seven samples were obtained from the outside vertical surface of two steel legs (Figure 4). Each sample was taken using a steel quadrat measuring 31.6 * 15.8 cm (0.05 m^2^) held in place by one of the two divers. The dimensions of the frame were identical to those used in previous studies (Coolen et al., 2018, 2015a). All marine growth within the frame was scraped off using a putty knife and the removed fauna was siphoned by an airlift sampler fed by a scuba tank. The airlift sampler was based on a combination of the samplers utilised in earlier reef studies (Coolen et al., 2018, 2015a). Modifications to the samplers described therein were as follows. The putty knife was attached to the airlift via a flexible ribbed hose (50 mm internal diameter). This was connected to a 48 mm (outer diameter) stainless steel pipe to which the air inlet was connected. The flow of air was regulated by a needle valve, which was connected to a scuba tank regulator by a 10-bar pressure hose. The stainless-steel pipe was connected to a straight 50 mm outer diameter pvc pipe ending in a 180° bend on top, made of 75 mm outer diameter PVC. This then ended into a sample net with a mesh size of 0.5 mm. The total length of the airlift above the air inlet was 150 cm. The screw cap nets, which were replaced for each sample, allowed an easy exchange during the dive. During a 50 minute dive, a two-person dive team was able to collect up to 12 samples.

**Figure 4.**
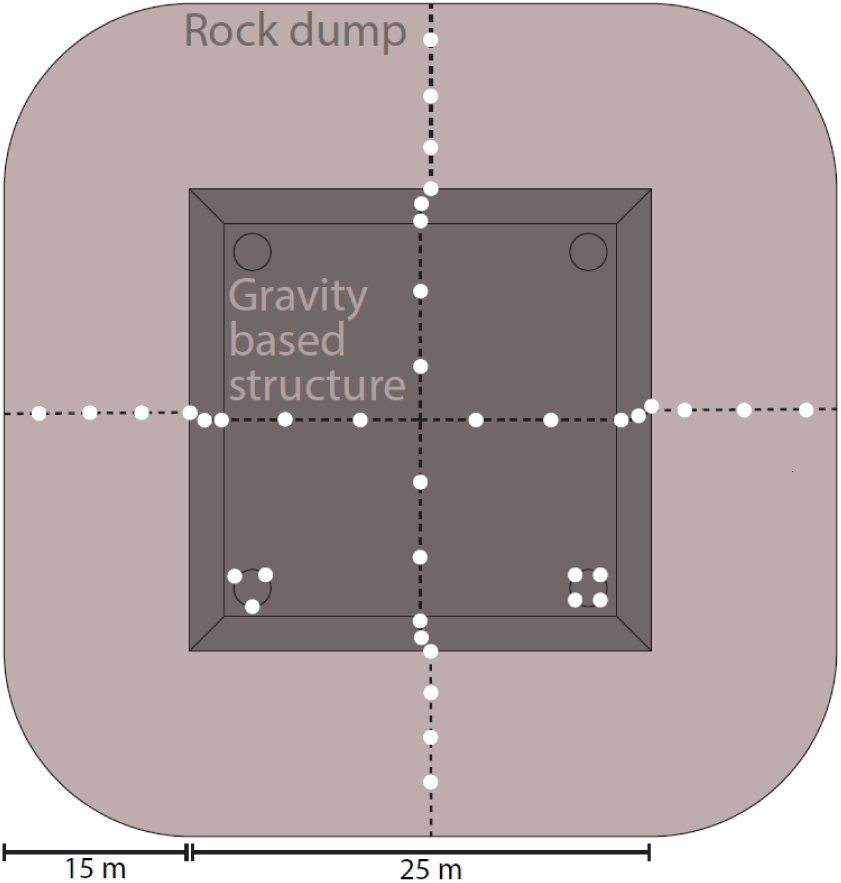
Sample locations. Schematic overview of the sampling design (top view, scale approximated). Samples (white dots) were taken in four directions on the gravity-based structure (GBS; dark colour), surrounding rock dump (light colour) and on legs (black rings in corners of GBS).

After collection, the samples were processed on board by depositing the collected macrofauna on a 212 µm mesh sieve, rinsing the nets with seawater to isolate all specimens. Ethanol (99%), measuring at least twice the volume of fauna, was then added to each sample. Within 48 hr, the ethanol was drained from each sample and replaced with fresh 99% ethanol, again with a volume of at least twice that of the fauna.

All samples were processed in the benthic laboratories of Bureau Waardenburg b.v. and Wageningen Marine Research. Each sample was sorted into major taxonomic groups, after which all specimens were identified to the lowest possible taxonomic level, mainly to species level. Where more than 200 individuals of the same species were present in a sample, species counts were undertaken by subsampling to a level where between 100 and 200 individuals of the species were left in a subsample.

After identification, the individuals from each non-colonial species from every sample with a wet weight >0.01 g were ash free dry weighed (6 hr at 500°C after drying) using a Precisa Gravimetrics prepASH 340 series. Colonial species such as Bryozoa, Hydrozoa, Porifera and Tunicata were not weighed or counted, but the area covered in a horizontal plane per species per sample was estimated to the nearest cm^2^ by flattening the species on grid paper.

#### 2.2.2. Seabed data

To allow a comparison of the macrofaunal assemblages present on the Halfweg GBS with those of its regional setting, seabed macrofaunal data were acquired from the published North Sea macrofauna dataset of the long term measurement programme MWTL (Monitoring Waterstaatkundige Toestand des Lands) in the Netherlands. The MWTL dataset is published with a creative commons zero licence by the Dutch Marine Information and Data Centre (Marine Information and Data Centre, 2019). It covers the entire Dutch part of the North Sea and includes data on macrofauna samples taken using various methods. For comparison with the airlifted macrofauna samples from Halfweg, only macrofauna data that were within range of 30 km from the Halfweg structure and were sampled using box corer and sieved on a 1 mm mesh (no smaller mesh size was available) were included. We appreciate that differences in sampling gear and mesh sizes used during sample processing with those from the GBS result in difficulties over direct comparisons with the data. Thus, caution will be applied when these results are compared with those of the GBS. The resulting dataset held 2,003 records from 118 samples. These samples had been taken between 1991 and 2015, with yearly samples between 1991 and 2010 as well as samples in 2012 and 2015.

#### 2.2.3. Artificial & natural reef data

A set of published marine growth data from reefs in the Dutch part of the North Sea (Coolen et al., 2018) was used to assess the uniqueness of the species on Halfweg on a larger scale. These data were acquired from scraped samples from five oil and gas structures, a wind farm and a rocky natural reef in the Borkum Reef Grounds. The locations are between 32 and 184 km distance from Halfweg. The distance from Halfweg to some of these locations, in particular to Borkum (164 km) and the D15-A platform (184 km) is large, but it is the most proximate dataset available that includes geogenic reef formations as well as the most detailed dataset on oil and gas platform fouling communities in the North Sea. The included artificial structures include ages both lower and higher than the 25 yr Halfweg has been in place, with an average age of 23 yr. No information on whether the installations had ever been treated with anti-fouling coatings was available. Although marine growth removal for inspection could have had an impact on the communities, previous analysis showed that whether an installation had been cleaned recently, had no significant effect on the species richness and only accounted for 0.3% of the variation (Coolen et al., 2018). Samples were taken by divers using a similar airlift as used for Halfweg (the platforms, rocky reef) or a sampling net (wind farm). The dataset from 145 samples contained only species abundances (individuals per m^2^ or presence-only for colonial species) and no biomass data.

#### 2.2.4. Data preparation

For data preparation and analysis, R version 3.6.1 (R Core Team, 2019) and Rstudio version 1.2.5001 (RStudio, 2019) were used. Prior to the analysis, all data were updated to include the most recent species names as published on the World Register of Marine Species (WoRMS Editorial Board, 2019), using the *wormsbynames* function from the worms package (Holstein, 2018). For the seabed data from the 30 km radius around Halfweg, seabed depth was obtained from EMODnet bathymetry data (EMODnet, 2019) using the extract function from the raster package (Hijmans, 2019). This resulted in a mean depth of 22 m from a range of 9 to 33 m. Mean depth of the Halfweg data was 21 m from a range of 17 to 24 m.

When samples included specimens that were not identified to species level, their abundance was added to a species in the lowest common higher taxon or removed when more than one species was present in the lowest common higher taxon in the same sample. Only individuals in samples with no species in a common higher taxon were left at the higher level (Coolen et al., 2018, 2015a). Since macrofaunal densities and weights in the seabed data were given per m^2^, all the marine growth data were converted to values per m^2^.

#### 2.2.5. Feeding traits

To provide a comparison of the feeding modes of the assemblages across the different datasets, the numerical composition of each assemblage across major feeding modes was assessed using biological traits. Each species was categorised across one or more of five feeding mode traits (suspension-feeders, deposit-feeders, predators, scavengers, parasites) using a fuzzy-coding approach based on the traits information used by Bolam et al. (2017, 2016). Fuzzy-coding allows the multi-faceted feeding behaviour of many species to be accounted for and overcomes the need to confine each species to a single mode of feeding. The feeding mode composition of each assemblage was calculated based on the most abundant species that, in total, accounted for >90% of the total abundance within each habitat.

#### 2.2.6. Data analysis

For each sample, species richness, Simpson biodiversity index (Simpson, 1949) using the diversity function from vegan package (Oksanen et al., 2019), total number of individuals and total ash free dry weight (AFDW) were calculated. Total observed and extrapolated species richness depends on the number of samples collected (Chao et al., 2014). The number of samples used to acquire the data for the seabed and other artificial structures was higher than for the concrete and rock dump of the GBS. Therefore, to compare total species richness among substrates, the mean total species richness was calculated from subsets of the seabed data and of artificial structures data. Subsets of 39 samples were randomly selected from each of these datasets. This process was repeated 10,000 times for both, and for every repetition the subset was used to calculate total species richness in all samples as well as extrapolated species richness based on the Chao estimate (Chao, 1987) using the specpool function (Oksanen et al., 2019). These numbers were also calculated for concrete, steel and the rock dump separately, for the combined Halfweg samples and for all available samples from the seabed, Borkum Reef Grounds and the other structures combined. A Euler plot (Euler, 1768) showing important overlap in species between substrates was generated using the euler function from the eulerr package (Larsson, 2019).

For each non-colonial species in the dataset, average abundance, AFDW biomass and standard errors were calculated for concrete, steel, rock dump and seabed. To test for differences in abundance and biomass between these substrates, generalised linear models were created, using a Gaussian distribution with log-link. The residuals were assumed to be normally distributed with a mean of 0 and variance of σ.

Community weighted means for all feeding traits in each substrate ∼ location combination were calculated using the weighted_mean function. Multivariate taxonomic structures were compared based on a Nonmetric Multidimensional Scaling (NMDS) plot created with the metaMDS function (Oksanen et al., 2019) using the Bray-Curtis dissimilarity index and 10,000 runs (Bray and Curtis, 1957). The ordination plot was used to visualise community differences as well as to assess multivariate spread to test whether the performance of permutational analysis of variance (PERMANOVA) was appropriate to test for community differences between the substrates (Anderson, 2005, 2001). A PERMANOVA (10,000 permutations, Bray–Curtis dissimilarity index) was performed to asses differences in community structure between substrates, using the adonis2 function (Oksanen et al., 2019).

Species status for the Netherlands (indigenous, non-indigenous, new observation) was assessed for all species observed on the Halfweg structures based on Bos et al. (2016) and the Dutch register of species (Naturalis and EIS, 2020) while the status of some of these species was updated based on a previous review (Coolen et al., 2018). Species not included in any of these publications were considered new species descriptions for the Dutch fauna.

## 3. RESULTS

All data underlying the results presented here are available as online supplement S2.

### 3.1 Species richness and uniqueness

In total, 65 species were observed on the Halfweg GBS and its associated rock dump. This included 52 species found on the concrete GBS, 44 species on the surrounding rock dump and 32 on the steel legs. Based on the extrapolated species richness, it was predicted that a total of 83 ± 11 (standard error) species are present on the combined structures of Halfweg, indicating that the survey approach resulted in an under-sampling of between 7 and 29 species (Table 1). Based on the iterative subsampling of 39 sand samples out of 118, the surrounding seabed exhibited 102 observed species, which were extrapolated to 151 ± 22. When all sand samples are included, the seabed comprised a total of 150 species, extrapolated to 265 ± 46 species. The other structures from Coolen et al. (2018) totalled 151 species, extrapolated to 178 ± 12. Simpson diversity index was highest on the rock dump at Halfweg (0.71 ± 0.03) but concrete (0.54 ± 0.06) was lower than seabed (0.69 ± 0.02). Lowest Simpson diversity was observed on the steel of other structures (0.34 ± 0.02).

**Table 1:**
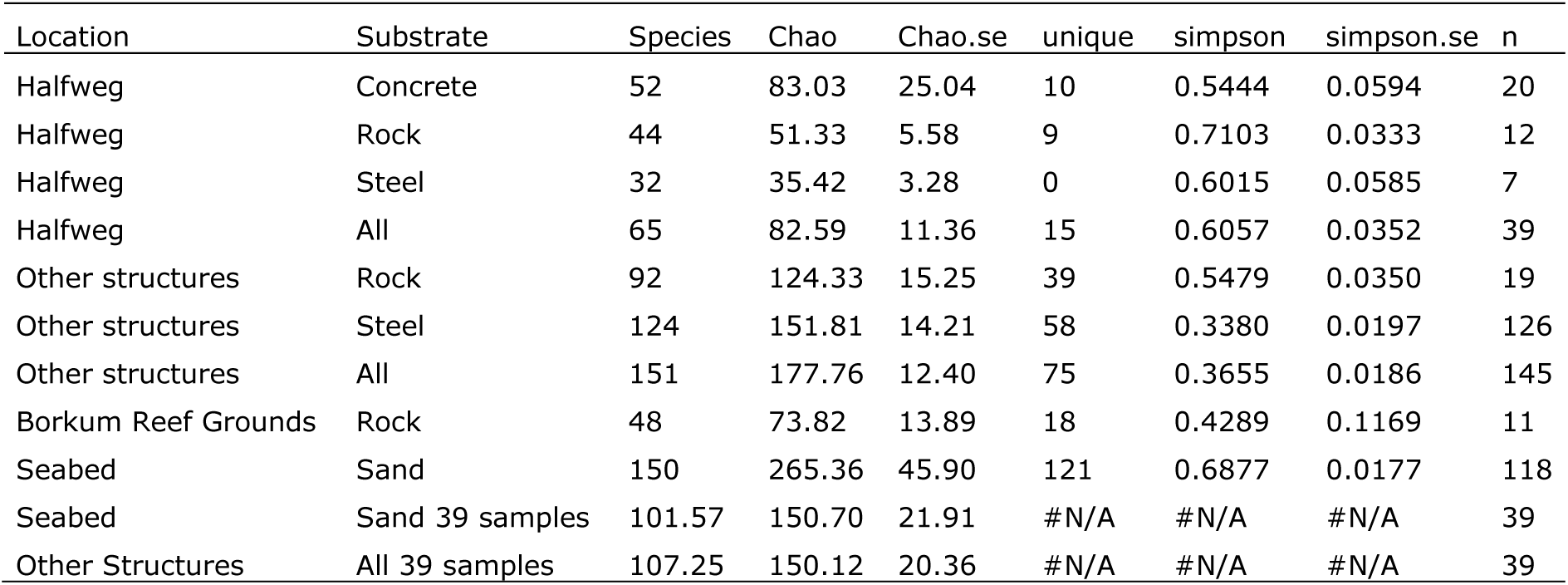
Species richness & uniqueness per substrate type. Location: the study sites, with ‘Other structures’ including oil and gas platforms and a wind farm. Substrate: the different types of substrates that were sampled where ‘all’ includes all substrates sampled at the location. Substrates include: Concrete: the GBS part of Halfweg, Rock: the rock dump part of Halfweg or other structures, Steel: the steel part of Halfweg or other structures. Species: total species richness, Chao: extrapolated species richness, Chao.se: standard error around the mean of extrapolated species richness, unique: number of species found only on that substrate (for concrete, rock & steel excluding samples from each other), simpson & simpson.se: mean Simpson diversity index with standard error, n: number of samples. Unique species were not calculated for the subsampling of seabed and structures.

Most substrate types revealed species that were not observed on any of the other substrates (Table 1). On the concrete of the GBS, 10 unique species were observed that were not observed on Borkum, other structures and the seabed. Six of them were also observed on the rock dump and steel, showing that the other four unique species on the concrete were not found on the rock dump, nor on steel. On rock dump, nine unique species were observed, out of which three species were not found on the GBS. The steel legs contained two additional unique species (out of four) that were not observed on the concrete and rocks. In total, 15 unique species occurred on the combined Halfweg substrates. Most overlap with the Halfweg substrates was observed with the other oil and gas and wind turbine structures. From the 52 species occupying the concrete structure, 30 (58%) were also observed on the rock dump around other structures and 36 (69%) on steel of other structures. From the 44 rock dump species, 27 (61%) occurred on the rock dump of other artificial structures and 31 (70%) on the steel of other structures. Highest relative overlap in species was observed between steel substrates, with 26 out of 32 species (81%) on the Halfweg steel observed on steel of other structures (Figure 5). The sandy seabed showed the lowest overlap with the Halfweg substrates, only 10 (7%) of the 150 seabed species were observed on the GBS, 6 (4%) on the rock dump and 2 (1%) on steel. When comparing natural and artificial hard substrates, the rocky substrates of the Borkum Reef Grounds had a relatively large overlap with Halfweg GBS, of 48 species, 12 (25%) were also observed on the GBS, another 12 (25%) on the rock dump and 10 (21%) on steel of Halfweg. Of a total of 311 species found in all available data, only seven (2%) were observed on six substrates (excluding seabed) and only the bryozoan *Electra pilosa* (<1%) on all seven substrates.

**Figure 5:**
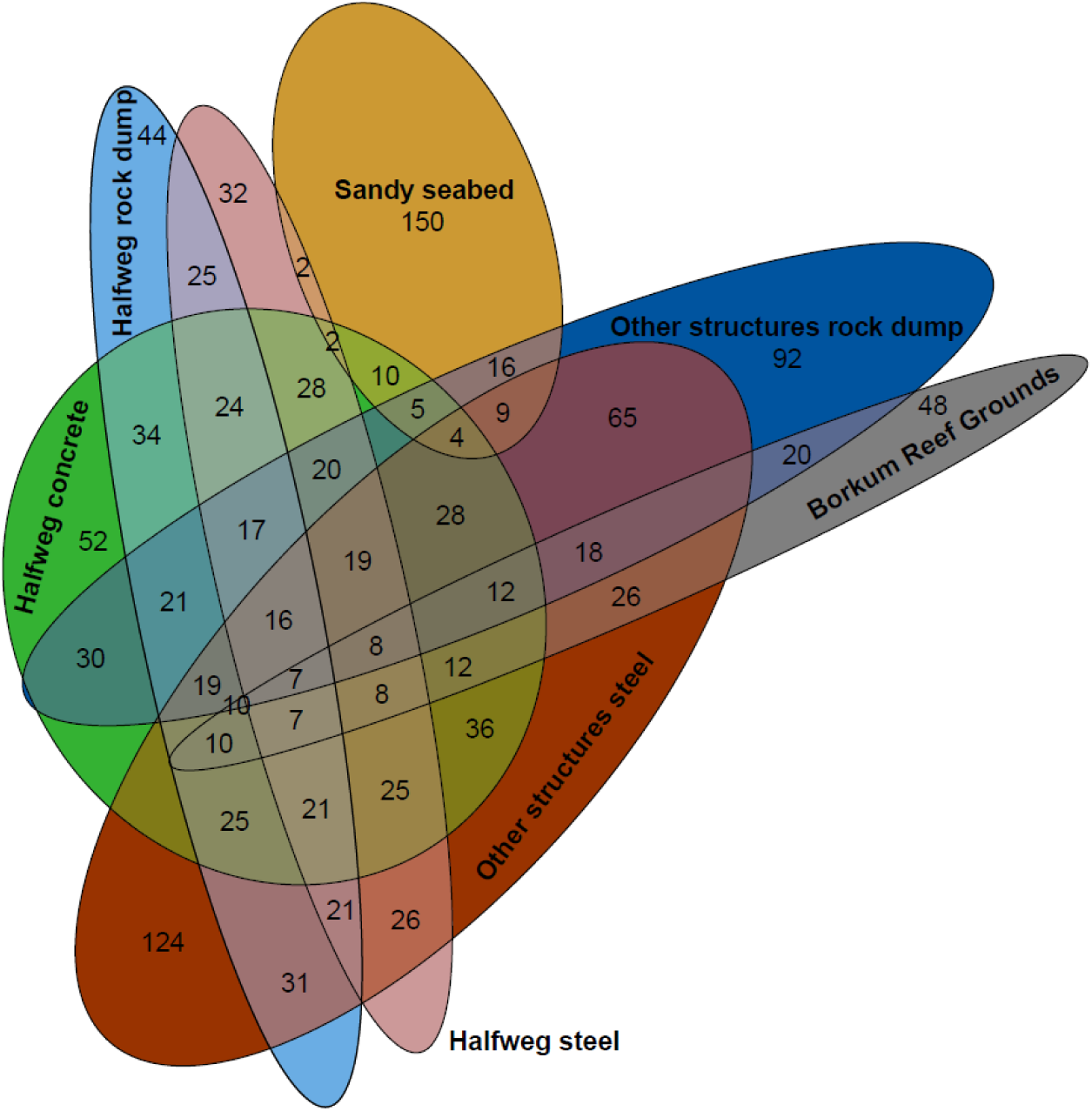
Euler diagram of overlap in species between substrates. Euler diagram presenting overlap in species between substrates (ellipses) for the strongest relations. Numbers present the total number of species per substrate (numbers in ellipse area without overlap) or shared between substrates (in the area of overlap between ellipses). Note that plotting all overlap between all combinations was impossible to visualise, resulting in missing overlap such as between the Sandy seabed and the Halfweg rock dump (6 species).

Two species (the amphipod *Monocorophium sextonae* and the colonial tunicate *Diplosoma listerianum*) found on Halfweg substrates were registered as non-indigenous for the Netherlands. Both species were also present on the other reefs. The bryozoan *Schizoporella unicornis* and the sponge *Leucosolenia botryoides* were not recorded in the species lists for the Netherlands (Bos et al., 2016; Naturalis and EIS, 2020) but *L. botryoides* was reported for the other structures (Coolen et al., 2018). *L. botryoides*, although not in the Dutch species list, has recently been reported from the Netherlands (Langeveld et al., 2020). It has originally been described from the English Channel (Ellis et al., 1786) and as such is here considered indigenous to the Netherlands. *S. unicornis* is an indigenous species in western Europe (Ryland et al., 2014).

### 3.2 Species abundance

Total abundance was significantly higher (p<0.01 R^2^=0.31) on the Halfweg substrates (mean abundance 40,393 ± 7,841 (standard error) per m^2^) than on the surrounding seabed (2,280 ± 165). Highest abundance was observed on concrete, with an average of 58,783 ± 13,739 individuals per m^2^. Arthropoda significantly accounted for the higher abundance on the Halfweg structure, with a mean abundance of 49,499 ± 10,341 individuals on concrete, 10,605 ± 2,276 on the rock dump and 25,080 ± 6,060 on steel (Figure 6). Within Arthropoda on the concrete, amphipods *Monocorophium acherusicum, Jassa herdmani, Phtisica marina* and *Stenothoe monoculoides* were responsible for 98% of the observed abundance. In the seabed only 654 ± 51 Arthropoda per m^2^ were observed, of which the amphipod *Urothoe poseidonis* provided 63%. In the seabed, Mollusca was the most abundant phylum with an average of 1,020 ± 163 individuals per m^2^. Nemertea were absent from all Halfweg and Borkum Reef Grounds samples. These were observed with a mean abundance of 44 ± 10 on the seabed and 18 on other platforms. Echinodermata were absent on the Halfweg rock dump and steel as well as on the geogenic rocky reefs of the Borkum Reef Grounds. On the Halfweg concrete 129 ± 86 Echinodermata individuals per m^2^ were observed, lower than the 466 ± 103 on other structures.

**Figure 6:**
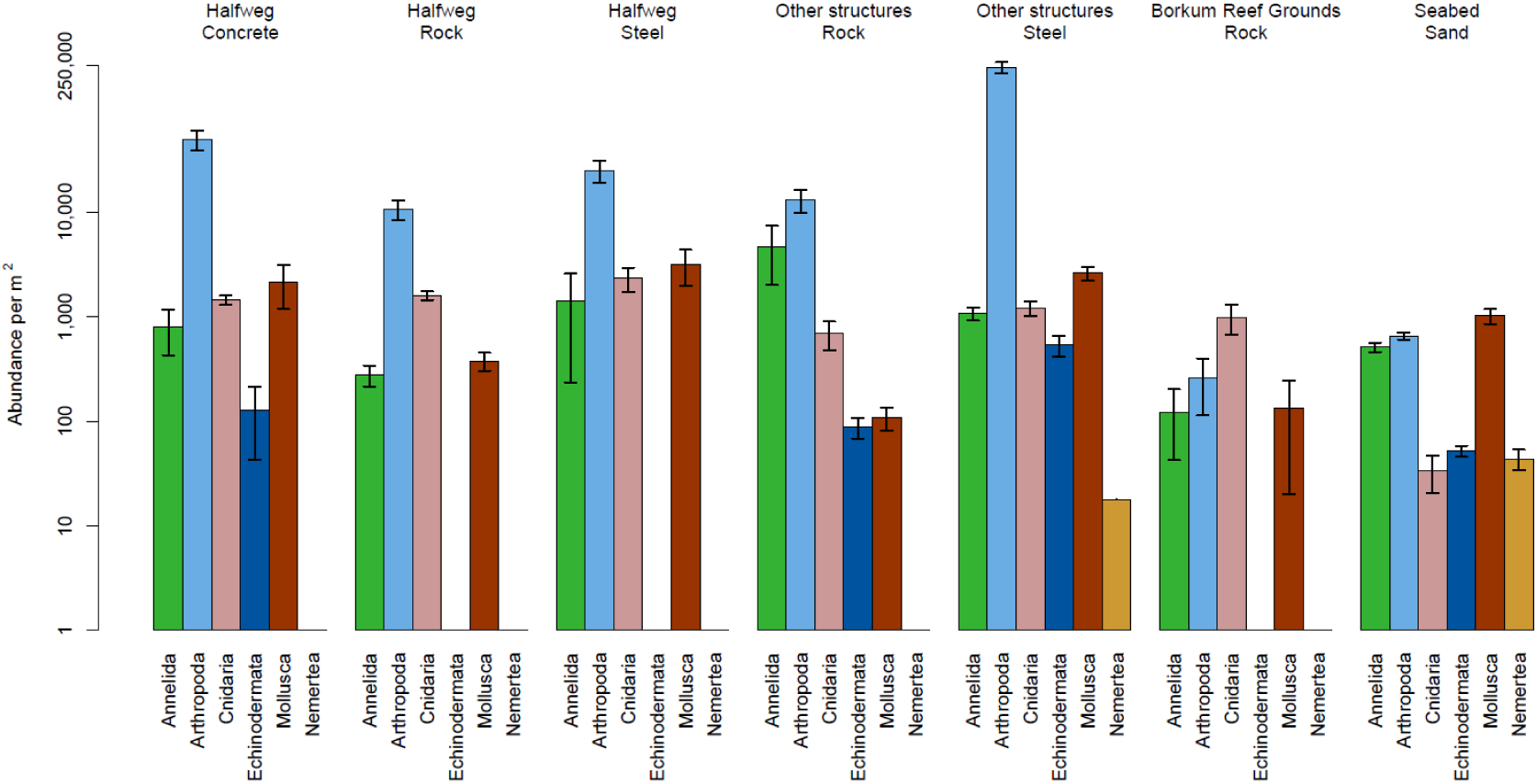
Average abundance. Bar plots showing average abundance with standard errors per m^2^ of non-colonial species per phylum on different substrates. Y-axis in log scale. Abundance of zero shown as empty slot with continued x-axis line.

### 3.3 Total biomass

Total biomass was significantly higher (p<0.001 R^2^=0.29) on the Halfweg substrates (mean AFDW of 204 g per m^2^ ± 20 standard error) than on the surrounding seabed (65 ± 8 g per m^2^). No significant difference of mean weight between the rock dump fauna (247 ± 37) and the concrete (191 ± 23) or steel (168 ± 65) was found (Table 2).

**Table 2:**
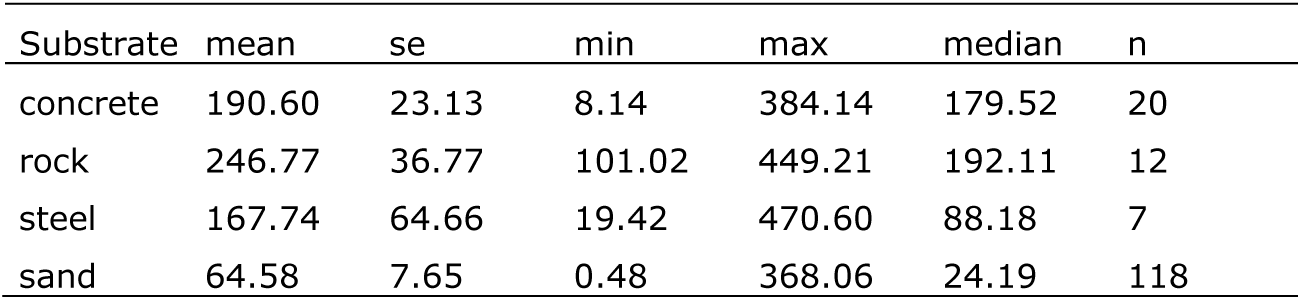
Ash free dry weights. Ash free dry weights (AFDW) in g per m^2^, with mean, standard error (se), minimum (min), maximum (max), median values and total number of samples (n). Concrete, Rock & Steel substrates based only on Halfweg samples.

| Substrate | mean | se | min | max | median | n |
| --- | --- | --- | --- | --- | --- | --- |
| concrete | 190.60 | 23.13 | 8.14 | 384.14 | 179.52 | 20 |
| rock | 246.77 | 36.77 | 101.02 | 449.21 | 192.11 | 12 |
| steel | 167.74 | 64.66 | 19.42 | 470.60 | 88.18 | 7 |
| sand | 64.58 | 7.65 | 0.48 | 368.06 | 24.19 | 118 |

The plumose anemone *Metridium senile* accounted for the high AFDW on concrete (183 (96%) ± 24 g AFDW per m^2^) and rock dump (245 (99%) ± 37 g AFDW per m^2^) of the GBS (Table 3). *M. senile* was also responsible for the high dominance of Cnidaria in most samples collected from the concrete and the rock dump (Figure 7). This species was not observed on the sandy seabed.

**Table 3.** Abundance and weight per species per location per habitat. Abundance of individual species in number (n) per m^2^, colonial species (with *) in cm^2^ per m^2^ on Halfweg and presence (P) on the other locations. Ash free dry weight in g per m^2^ (no data for colonial species [marked with *], *Balanus crenatus* and *Verruca stroemia*) of species observed on Halfweg and the seabed. All numbers ± standard error.

| Phylum / species | Halfweg<br>Concrete<br>n or cm2* | Halfweg<br>Concrete<br>AFDW | Halfweg<br>Rock<br>n or cm2* | Halfweg<br>Rock<br>AFDW | Halfweg<br>Steel<br>n or cm2* | Halfweg<br>Steel<br>AFDW | Seabed<br>Sand<br>n | Seabed<br>Sand<br>AFDW | Other structures<br>Steel<br>n | Other structures<br>Rock<br>n | Borkum<br>Rock<br>n |
| --- | --- | --- | --- | --- | --- | --- | --- | --- | --- | --- | --- |
| **Annelida** |  |  |  |  |  |  |  |  |  |  |  |
| *Aonides paucibranchiata* | 1±1 | 0 | 0 | 0 | 0 | 0 | 0 | 0 | 0 | 0 | 0 |
| *Dipolydora coeca* | 3±1.64 | 0 | 6.67±6.67 | 0 | 0 | 0 | 0 | 0 | 0 | 0 | 0 |
| *Eulalia viridis* | 1±1 | 0.01±0.01 | 8.33±6.72 | 0.01±0.01 | 5.71±5.71 | 0 | 0 | 0 | 111.51±24.69 | 27.41±18.99 | 14.55±14.55 |
| *Eunereis longissima* | 18±6.94 | 0.01±0.01 | 5±3.59 | 0 | 0 | 0 | 2.14±0.71 | 0.35±0.17 | 35.91±20.63 | 84.21±53.87 | 3.64±2.44 |
| *Harmothoe impar* | 3±1.64 | 0 | 0 | 0 | 0 | 0 | 0 | 0 | 50.54±17.2 | 2.82±2.05 | 1.82±1.82 |
| *Lepidonotus squamatus* | 8±6.05 | 0.01±0.01 | 0 | 0 | 5.71±5.71 | 0.01±0.01 | 0 | 0 | 60.71±9.67 | 30.15±17.1 | 5.45±2.82 |
| *Mysta picta* | 0 | 0 | 1.67±1.67 | 0.01±0.01 | 0 | 0 | 0 | 0 | 0 | 0 | 0 |
| *Phyllodoce mucosa* | 2±2 | 0 | 3.33±3.33 | 0 | 0 | 0 | 8.04±7.1 | 0.04±0.04 | 1.56±0.64 | 0 | 0 |
| *Sabellaria spinulosa* | 194±79.75 | 0.32±0.09 | 141.67±46.41 | 0.24±0.05 | 0 | 0 | 0 | 0 | 1.98±1.07 | 3658.61±2238.14 | 3.64±3.64 |
| *Syllis gracilis* | 571±295 | 0.1±0.07 | 111.67±28.44 | 0.01±0 | 1240±1113.18 | 0.12±0.12 | 0 | 0 | 22.86±10.76 | 0 | 0 |
| *Syllis vittata* | 0 | 0 | 0 | 0 | 165.71±97.27 | 0.01±0.01 | 0 | 0 | 0 | 0 | 0 |
| **Arthropoda** |  |  |  |  |  |  |  |  |  |  |  |
| *Aora gracilis* | 0 | 0 | 56.67±35.75 | 0 | 5.71±5.71 | 0 | 0 | 0 | 0 | 7.37±5.12 | 47.27±45.31 |
| *Apolochus neapolitanus* | 0 | 0 | 5±3.59 | 0 | 0 | 0 | 0 | 0 | 0 | 0 | 0 |
| *Balanus crenatus* | 45±20.36 | 0 | 65±24.14 | 0 | 5.71±3.69 | 0 | 0.43±0.43 | 0 | 32.69±9.57 | 71.13±67.2 | 0 |
| *Bodotria scorpioides* | 3±3 | 0 | 1.67±1.67 | 0 | 0 | 0 | 0 | 0 | 0 | 0 | 0 |
| *Cancer pagurus* | 11±5.71 | 0.06±0.04 | 3.33±2.25 | 0.03±0.03 | 17.14±6.8 | 0.3±0.22 | 0 | 0 | 5.79±1.17 | 1.05±1.05 | 0 |
| *Cumopsis goodsir* | 0 | 0 | 0 | 0 | 2.86±2.86 | 0 | 0 | 0 | 0 | 0 | 0 |
| *Gitana sarsi* | 125±65.11 | 0 | 85±17.08 | 0 | 151.43±89 | 0 | 0 | 0 | 143.58±37.49 | 0 | 0 |
| *Jassa herdmanni* | 8536±2384.97 | 1.8±0.57 | 1580±465.21 | 0.14±0.05 | 19691.43±5117.24 | 6.92±2.34 | 0 | 0 | 204173.34±27866.63 | 8090.9±2981.26 | 0 |
| *Melita palmata* | 1±1 | 0 | 0 | 0 | 0 | 0 | 0 | 0 | 0 | 0 | 0 |
| *Microprotopus maculatus* | 1±1 | 0 | 8.33±5.2 | 0 | 0 | 0 | 0.12±0.12 | 0 | 0 | 601.05±332.61 | 0 |
| *Monocorophium acherusicum* | 35667±8207.31 | 4.07±0.96 | 6348.33±1935.14 | 0.8±0.25 | 694.29±264.38 | 0.01±0.01 | 0 | 0 | 437.43±200.41 | 1653.83±477.1 | 43.64±24.69 |
| *Monocorophium sextonae* | 19±16.12 | 0 | 0 | 0 | 0 | 0 | 0 | 0 | 469.66±213.12 | 177.29±75.06 | 0 |
| *Phthisica marina* | 2981±584.35 | 0.32±0.04 | 2081.67±305.44 | 0.35±0.05 | 1808.57±849.95 | 0.17±0.09 | 0 | 0 | 619.82±143.26 | 66.43±18.75 | 0 |
| *Pilumnus hirtellus* | 66±39.34 | 0.02±0.01 | 6.67±3.76 | 0 | 105.71±48.93 | 0.03±0.02 | 0 | 0 | 27.44±5.68 | 2.82±2.05 | 0 |
| *Pisidia longicornis* | 156±85.15 | 0.09±0.03 | 33.33±21.65 | 0.07±0.06 | 51.43±28.24 | 0.06±0.03 | 0 | 0 | 63.38±15.61 | 1148.23±649.34 | 0 |
| *Pseudocuma simile* | 1±1 | 0 | 0 | 0 | 0 | 0 | 0.98±0.38 | 0 | 0 | 0 | 0 |
| *Stenothoe monoculoides* | 1520±292.45 | 0.05±0.01 | 293.33±64.82 | 0 | 2537.14±782.25 | 0.06±0.04 | 0 | 0 | 3214.14±638.4 | 342.71±119.17 | 0 |
| *Stenothoe valida* | 1±1 | 0 | 0 | 0 | 0 | 0 | 0 | 0 | 423.71±101.36 | 13.16±10.56 | 0 |
| *Verruca stroemia* | 366±228 | 0 | 36.67±19.82 | 0 | 8.57±4.04 | 0 | 0 | 0 | 0.6±0.3 | 255.75±227.43 | 0 |
| **Bryozoa** |  |  |  |  |  |  |  |  |  |  |  |
| *Alcyonidioides mytili\** | 0 | 0 | 25±21.62 | 0 | 0 | 0 | 0 | 0 | P | P | 0 |
| *Arachnidium fibrosum\** | 0 | 0 | 38.33±36.55 | 0 | 0 | 0 | 0 | 0 | P | P | 0 |
| *Bugulina turbinata\** | 570±478.42 | 0 | 6.67±2.84 | 0 | 62.86±56.3 | 0 | 0 | 0 | 0 | P | 0 |
| *Callopora dumerilii\** | 0 | 0 | 25±25 | 0 | 0 | 0 | 0 | 0 | P | P | 0 |
| *Conopeum reticulatum\** | 4±2.34 | 0 | 331.67±329.85 | 0 | 0 | 0 | 0 | 0 | P | P | P |
| *Crisularia plumosa\** | 172±159.51 | 0 | 13.33±9.95 | 0 | 0 | 0 | 0 | 0 | 0 | 0 | 0 |
| *Electra pilosa\** | 41±31.71 | 0 | 20±9.53 | 0 | 8.57±4.04 | 0 | P | 0 | P | P | P |
| *Microporella ciliata\** | 0 | 0 | 1.67±1.67 | 0 | 0 | 0 | 0 | 0 | P | P | 0 |
| *Schizomavella linearis\** | 72±37.12 | 0 | 0 | 0 | 57.14±47.89 | 0 | 0 | 0 | P | P | 0 |
| *Schizoporella unicornis\** | 5±3.52 | 0 | 11.67±11.67 | 0 | 8.57±5.95 | 0 | 0 | 0 | 0 | 0 | 0 |
| *Scruparia chelata\** | 32±8.13 | 0 | 6.67±2.84 | 0 | 5.71±3.69 | 0 | 0 | 0 | P | 0 | 0 |
| *Scrupocellaria scruposa\** | 547±510.41 | 0 | 26.67±9.64 | 0 | 0 | 0 | 0 | 0 | 0 | 0 | 0 |
| **Chordata** |  |  |  |  |  |  |  |  |  |  |  |
| *Diplosoma listerianum\** | 601±542.61 | 0 | 28.33±7.57 | 0 | 651.43±368.96 | 0 | 0 | 0 | P | P | P |
| **Cnidaria** |  |  |  |  |  |  |  |  |  |  |  |
| *Clytia hemisphaerica* * | 57±31.76 |  | 8.33±4.58 |  | 5.71±3.69 |  | 0 |  | P | P | P |
| *Ectopleura larynx* * | 0 |  | 0 |  | 42.86±42.86 |  | 0 |  | P | P | P |
| *Laomedea flexuosa* * | 0 |  | 3.33±3.33 |  | 0 |  | 0 |  | P | 0 | 0 |
| *Metridium senile* | 1361±155.04 | 183.4±23.67 | 1580±152.53 | 245.05±36.82 | 1948.57±586.95 | 157.56±66.92 | 0 | 0 | 715.97±130.66 | 594.21±220.44 | 987.27±317.77 |
| *Obelia bidentata* * | 0 |  | 8.33±8.33 |  | 0 |  | 0 |  | P | P | P |
| *Plumularia setacea* * | 33±18.08 |  | 1.67±1.67 |  | 5.71±3.69 |  | 0 |  | 0 | 0 | 0 |
| Echinodermata |  |  |  |  |  |  |  |  |  |  |  |
| *Amphipholis squamata* | 5±4.07 | 0 | 0 | 0 | 0 | 0 | 0 | 0 | 102.22±54.8 | 0 | 0 |
| *Ophiorthrix fragilis* | 36±31.87 | 0 | 0 | 0 | 0 | 0 | 0 | 0 | 91.49±46.08 | 7.37±4.64 | 0 |
| *Ophiura albida* | 1±1 | 0 | 0 | 0 | 0 | 0 | 2.43±0.68 | 0.01±0.01 | 0 | 0 | 0 |
| *Ophiura ophiura* | 3±3 | 0 | 0 | 0 | 0 | 0 | 1.53±0.78 | 0.06±0.03 | 0 | 0 | 0 |
| Mollusca |  |  |  |  |  |  |  |  |  |  |  |
| *Brachystomia scalaris* | 19±19 | 0 | 0 | 0 | 0 | 0 | 0 | 0 | 389.43±121.53 | 2.82±2.82 | 0 |
| *Epitonium clathratulum* | 363±82.54 | 0.01±0.01 | 273.33±68.37 | 0.01±0.01 | 8.57±5.95 | 0 | 0 | 0 | 0.14±0.14 | 0.94±0.94 | 0 |
| *Epitonium clathrus* | 0 | 0 | 1.67±1.67 | 0 | 0 | 0 | 0 | 0 | 0 | 0 | 0 |
| *Euspira nitida* | 56±20.45 | 0 | 3.33±2.25 | 0 | 0 | 0 | 9.68±1.69 | 0.1±0.04 | 0 | 0 | 0 |
| *Hiatella arctica* | 1±1 | 0 | 0 | 0 | 5.71±5.71 | 0 | 0 | 0 | 3.81±2.12 | 1.88±1.29 | 0 |
| *Kellia suborbicularis* | 34±13.07 | 0 | 1.67±1.67 | 0 | 62.86±62.86 | 0.02±0.02 | 0 | 0 | 6.67±3.99 | 0 | 0 |
| *Mytilus edulis* | 1509±930.33 | 0.18±0.11 | 98.33±71.16 | 0.03±0.02 | 3100±1146.89 | 1.02±0.37 | 0 | 0 | 1728.83±322 | 32.22±15.34 | 3.64±3.64 |
| *Pusillina inconspicua* | 67±32.55 | 0 | 0 | 0 | 0 | 0 | 0 | 0 | 0 | 0 | 0 |
| *Spiralinella spiralis* | 1±1 | 0 | 0 | 0 | 0 | 0 | 0 | 0 | 0 | 0 | 0 |
| *Venerupis corrugata* | 1±1 | 0 | 0 | 0 | 0 | 0 | 0.11±0.11 | 0.01±0.01 | 0.85±0.4 | 0.94±0.94 | 0 |
| Porifera |  |  |  |  |  |  |  |  |  |  |  |
| *Halichondria panicea* * | 1362±885.94 |  | 121.67±78.72 |  | 334.29±330.96 |  | 0 |  | P | 0 | 0 |
| *Leucosolenia botryoides* * | 35±31.88 |  | 0 |  | 2.86±2.86 |  | 0 |  | P | 0 | 0 |

**Figure 7:**
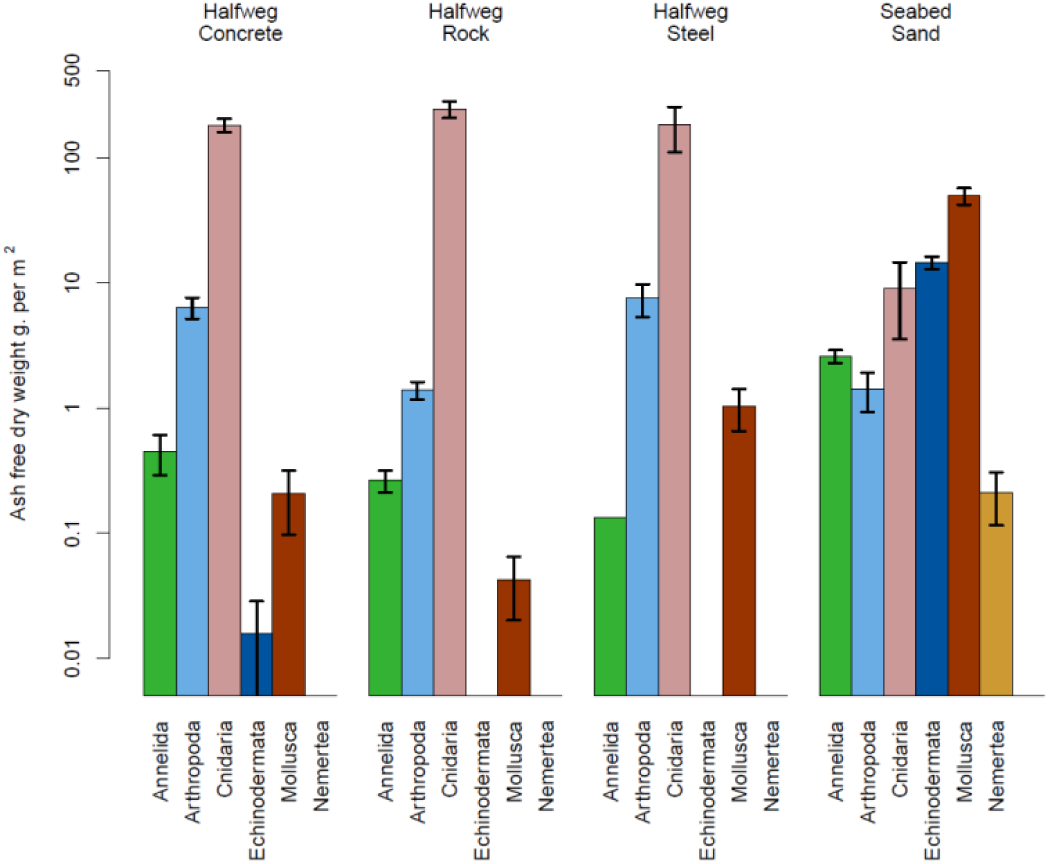
Average total biomass. Bar plots showing average ash free dry weights with standard errors per m^2^ of non-colonial species per phylum on concrete, steel, rock dump and the sandy seabed. Y-axis in log scale. No data were available for the Borkum Reef Grounds and other structures.

The biomass of the surrounding seabed was dominated by Mollusca (50 g AFDW per m^2^). The subtruncate surf clam *Spisula subtruncata* was the dominating species, with an average AFDW of 35 (54% of total biomass) ± 7 g per m^2^. This species was absent on Halfweg, which exhibited a total Mollusca biomass below 1 g AFDW per m^2^.

The total biomass on the concrete GBS was estimated to be 262 kg AFDW (based on the average weight times the available substrate), 20 kg on the steel legs and 2,564 kg on the surrounding rock dump. Therefore, the total weight of the macrofauna on the Halfweg substrates was 2,846 kg. The area of seabed covered by the GBS by both hard substrates was estimated to be 3,617 m^2^ which, based on the mean total biomass per m^2^ in the seabed, would hold a total macrofauna weight of 232 kg if Halfweg was absent.

### 3.4 Feeding traits

The feeding trait analysis indicated similarities between the concrete and rock dump of Halfweg and the seabed, where deposit feeding was similar (rock and seabed) or higher (concrete) to suspension feeding (Figure 8). On all other hard substrates, including the steel legs of Halfweg, suspension feeding was the main feeding trait, followed by scavenging and predation, while deposit feeders were very few or even absent (other steel structures).

**Figure 8:**
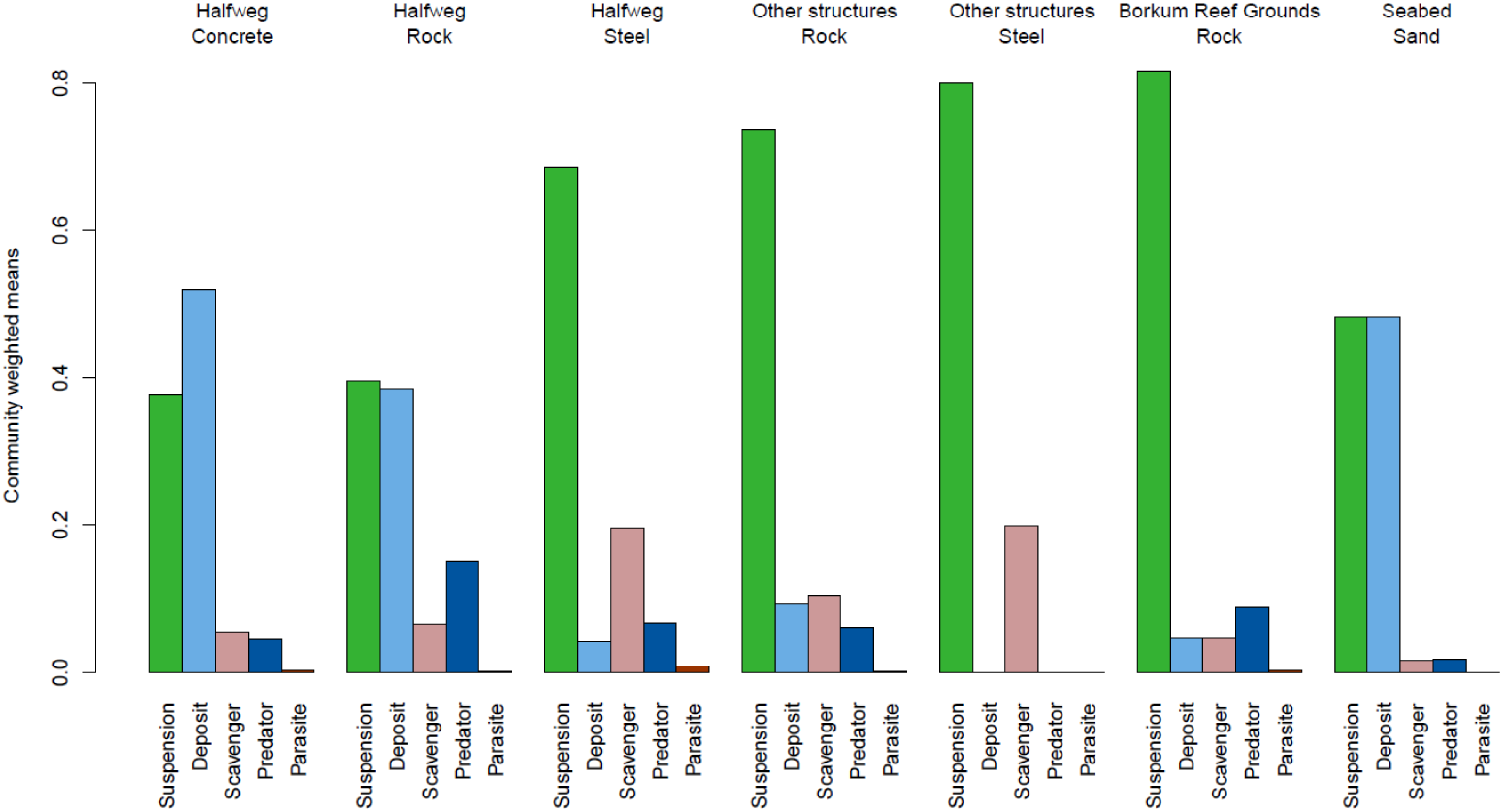
Feeding traits per structure. The feeding traits observed in the different structures under study. The traits are expressed as community weighted means.

### 3.5 Community differences

The non-metric multidimensional scaling (NMDS; 2 dimensions, stress 0.11; Figure 9) of all available abundance data showed a cluster of all reef samples which was clearly separated from those of sandy seabed samples. Within the cluster of reef substrates, the Halfweg substrates cluster together, although some overlap with those from the other structures is also evident. Samples taken from the Halfweg concrete and steel show comparatively less variation than those of other substrates. The assemblages of the samples from the Borkum Grounds rocks appear very variable, with two being located at the top of the plot and the remainder at the bottom. PERMANOVA showed a significant (p<0.001) influence of substrate type on community structure, confirming the observed differences observed in the NMDS.

**Figure 9:**
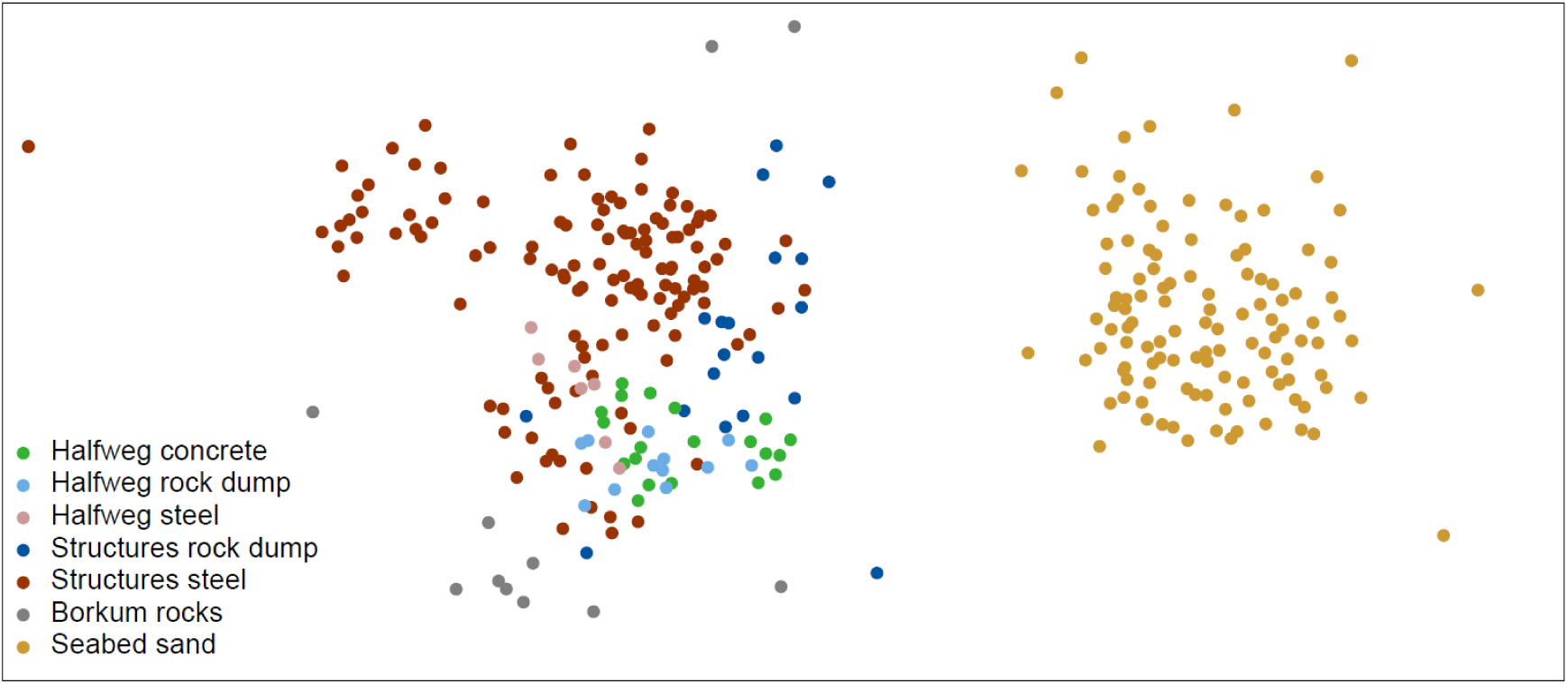
Non-metric multidimensional scaling plot. NMDS plot based on abundance data from all acquired data (Bray-Curtis dissimilarity index, 10,000 runs, stress 0.11, best solution after no convergence).

## 4. DISCUSSION

### 4.1 Ecological importance of the Halfweg GBS structure

The presence of the Halfweg structure has a notable effect on both the local biodiversity and structuring of the macrofauna community. While diversity comparisons between hard substratum habitats and sedimentary regions must be made with caution due to differences in the manner in which the faunal assemblages of these fundamentally different substrata are assessed, a total of 65 species were observed on the relatively limited spatial extent of the Halfweg structure, while a total of 150 species were observed in the surrounding seabeds within a 30 km radius. Only 10 of the Halfweg species were also observed in the seabed. The presence of the artificial and rocky substrates of the Halfweg GBS, therefore, increased local species richness by 55 (37% compared to 118 seabed samples, 53% when compared to 39 seabed samples). On the complete Halfweg structure, totalling 3,617 m^2^, the macrofauna AFDW biomass was 12 times that of the comparable area of sandy seabed. This increase results from a combination of a higher biomass per m^2^ of area available to fouling species and from the three-dimensional structural complexity of the concrete and rock dump increasing the available area compared to a relatively flat sedimentary seabed (Calow, 1972; Cooper and Testa, 2001; Graham et al., 1988).

The high Simpson diversity observed on the Halfweg rock dump and comparatively low diversity on the concrete is in harmony with observations at wind farms where straight steel monopiles have been demonstrated to harbour a lower diversity than surrounding rocky natural substrates (Wilhelmsson and Malm, 2008). A comparable ratio was also found on the steel vs the rock dump around the other structures assessed here. It is noteworthy that mean Simpson diversity at the Borkum reef grounds was lower than all other artificial structures in this study, which may have been caused by under-sampling, as suggested by the large standard error around Borkum reef grounds diversity.

Our data revealed that 23% of the species observed on Halfweg were unique and were not observed on other platforms, on wind turbines foundations nor a natural reef between 32 and 184 km from Halfweg. When only considering the concrete GBS, 10% of the species was not found on any of the other substrates, rising to 19% when excluding the Halfweg rock dump and steel from this comparison. On the rock dump a very similar percentage of 9 was not found elsewhere, 21% when excluding the GBS and steel. However, such localised species distributions may not be unusual: all other reefs from the data in this study also showed a high number of unique species. Sampling such habitats is logistically challenging and the resulting datasets, including those used here (Coolen et al., 2020, 2018, 2016b, 2015a), are often far from comprehensive. The Chao species richness estimator, when applied to these data, revealed that such sampling does result in underestimations of the species present on such structures. Indeed, for the Halfweg GBS, the Chao species richness estimates suggest that observed richness on the concrete of the GBS might double based on the acquisition of a sufficient number of samples. Thus, uniqueness of species on the GBS and on the other artificial substrates may have been over-estimated in this study as a consequence of limited sampling. However, this study clearly indicates that the Halfweg structure increases the local species richness, suggesting positive impacts of this artificial reef on the macrofaunal biodiversity. This has been confirmed for other artificial habitats in the North Sea, such as ship wrecks (Coolen et al., 2015a; Zintzen, 2007), offshore wind turbines (De Mesel et al., 2015) and oil and gas platforms (Coolen et al., 2018). Enhanced macrofaunal biodiversity on artificial reefs is considered to be a beneficial ecosystem effect (Causon and Gill, 2018).

Two non-indigenous species (3% of the total) were observed on Halfweg, which is similar to the percentage reported from the deep parts of the other structures by Coolen et al. (2018). Both species, the amphipod *Monocorophium sextonae* and the colonial tunicate *Diplosoma listerianum*, have previously been reported from either natural or artificial reef structures in the North Sea (Coolen et al., 2018, 2015a; De Mesel et al., 2015; Vance et al., 2009; Zintzen et al., 2007; Zintzen and Massin, 2010) and can be considered common on reefs in the North Sea. Several authors have suggested that offshore infrastructure is likely to play a role in the distribution of non-indigenous species (Adams et al., 2014; IPIECA, 2010; Macreadie et al., 2011). The capacity of such structures to act as stepping stones, whereby species’ distributions are extended, depends on their depth (Coolen et al., 2016b) together with species’ life history strategies (Coolen et al., 2020). The bryozoan *Schizoporella unicornis*, found on the concrete, steel and rock dump of the GBS, is considered native to western Europe (Ryland et al., 2014) but was not previously reported from within the marine waters of the Netherlands. It is possible that the species has always been present in the Netherlands but missed in previous surveys as has been suggested for first observations of other species on artificial structures in the North Sea (Coolen et al., 2015b; Dias et al., 2017; Faasse et al., 2016).

Within Halfweg, feeding functionality differs among substrates. Most noteworthy is the dominance of suspension feeders on the steel of the GBS which is similar to other steel structures (platform jackets and wind turbine foundations), compared to an increased dominance of deposit feeders on the concrete and rock on the GBS which was more in line with assemblages of the surrounding seabed. The similarities between the concrete and the seabed, and the evident differences between Halfweg concrete and the other artificial hard substrates may result from a number of factors. The similarities with the soft seabed could be the result of the combination of different deposit feeding mechanisms, i.e. surface and subsurface deposit feeders, into one general deposit feeding trait. Separating the deposit feeding mechanisms into these two categories could have resulted in more differentiation between these two habitats, since there is a lack of sub-surface deposit feeders on hard substrates and since the most abundant arthropod on the seabed, the burrowing amphipod species *Urothoe poseidonis*, is a sub-surface deposit feeder (Kröncke et al., 2013). The high relative abundance of deposit feeding organisms on the Halfweg concrete compared to the other artificial structures could be the after-effect of the colonisation of the jacket foundation that was on top of this concrete substrate for a prolonged period which was removed eight months before sampling. Fouling organisms inhabiting vertical hard substrates such as oil and gas platforms and offshore wind turbines produce biodeposits that sink on the seabed, causing very local accumulation of organic matter (Coates et al., 2014). The increased local organic matter accumulation provides resources for multiple species, increasing the local trophic diversity and attracting species with distinct feeding habits (Mavraki et al., 2020a). Therefore, organisms with different feeding traits are to be expected at the base (i.e. Halfweg concrete and rock dump) of such structures. It is, however, noteworthy that the rock dump around other structures did not show these high numbers of deposit feeders. This may have been caused by differences in environmental conditions between locations which have not been included in our analysis. Finally, the orientation of the substrates could have had an effect on the feeding traits observed for the different types of structures resulting in clear differences between horizontally oriented substrates (part of the Halfweg concrete and seabed) and vertical structures (side walls of Halfweg, its legs and the other steel structures). Studies have indicated that substrate orientation has a large effect on community composition (Boström et al., 2010; Glasby and Connell, 2001), which could explain the variations observed between horizontally and vertically oriented habitats in terms of feeding traits.

The current study has limitations that should be noted while interpreting the results. The most proximate other structure used for comparisons with Halfweg was 32 km away, while the most distant structure was at the Cleaver Bank, 184 km from Halfweg. The Borkum Reef Grounds were 164 km from Halfweg. The species communities on the distant reefs may have differed from Halfweg due to different environmental conditions. The more distant, offshore reefs are located in clearer water with lower chlorophyll a concentrations, lower summer temperatures and lower water currents (Coolen et al., 2016a; van der Stap et al., 2016). However, several of the structures, including those at Borkum Reef, were located within 50 km from the coast, therefore minimising the effect of these environmental differences on the faunal data. The only other geogenic reef that is closer to Halfweg is Texel Rough (at 50 km; Coolen et al., 2018). However, no data on fouling communities are available from Texel Rough. Therefore the comparison with the Borkum Reef Grounds was the only available option in this study. Although the average installation age was similar to Halfweg, all the structures included in the comparison were of different age than Halfweg. However, previous analyses of these other data have demonstrated a limited effect of age on the macrofaunal assemblages compared to environmental or biotic effects. Within year variation was suggested to be much more important than long term yearly variation (Coolen et al., 2018).

### 4.2 Removal options

Two removal options are currently being considered: the GBS is fully removed and 50% of the rock dump is scattered across the area where the GBS was present while the remaining rock dump is left untouched, and; the GBS is left in place and all the rock dump remains untouched.

The data acquired during this study allow a simplistic comparison of the implications of the two options on localised macrofaunal diversity to be undertaken. In the first scenario, all the fauna that is currently present on the concrete GBS will be removed or scattered during removal, in essence resulting in the removal of most of the species present on the concrete. An estimated 50% of the rock dump will be placed elsewhere or dredged up (personal communication Gert de Raadt, Petrogas). When moved by crane, some of the macrofauna may remain on the rock dump, although it is unlikely that the rocks will be placed in an identical orientation, likely resulting in the burial or damaging of most fauna present on the moved rock dump. Assuming 50% of the rock will remain untouched, the fauna present on these rocks will mostly remain. On a longer term, the moved rock dump is likely to be colonised by a similar community as is currently present, originating from the untouched rock dump, small hard substrates on the seabed or other source locations where the current species originate from, e.g. by colonisation by settling larvae or via migration of juveniles and adults (Coolen et al., 2020; Krone and Schröder, 2011; Luttikhuizen et al., 2019). In this instance, species richness may recover to approximately the 44 species that were observed on the rock dump in this study. This scenario will therefore result in the loss of 21 macrofaunal species that are currently present on the concrete and steel but not on the rock dump. However, four of these 21 species have also been observed in the seabed, so the net loss of species richness to the local area is predicted to be 17 species or more as the observed species richness is, as we have demonstrated, not exhaustive. No unique species could be calculated for extrapolated species richness but given that these numbers are higher than observed richness the net loss might be higher than 17. Since after the removal of the GBS, the rock dump will be scattered across the area, the total area of substrate available to fouling communities could remain approximately like what is presently available on the GBS plus rock dump. Assuming this would be similarly colonised, this may result in a total biomass comparable to the current biomass. Feeding mode diversity is likely to be reduced when Halfweg GBS is removed, as the communities on the steel parts of the structure show different dominant feeding traits than present on the seabed. This would lead to a shift towards suspension feeding, reducing the functional evenness since species share a specific functional trait (Mason et al., 2005).

In the second scenario, the lack of management intervention will results in no species or biomass loss. The GBS and the rock dump have been in place since 1996 and have possibly reached, although this was not possible to assess in this study, a stable stage of succession (Oshurkov, 1992). Observations from other structures, which were both older and younger than the Halfweg GBS, support this as each has revealed a *Metridium senile* dominated community at depths similar to Halfweg (Coolen et al., 2018; Krone et al., 2013; Whomersley and Picken, 2003).

## 5. CONCLUSION

The presence of Halfweg, including its different materials in the form of steel, concrete and rock, has a clear effect by increasing local species richness as well as feeding mode diversity. Removal of the gravity-based foundation of Halfweg will result in the loss of a significant number of species from the local area and possibly a lowering of feeding mode diversity. Due to the scattering of the rock dump after removal, local impact on macrofouling biomass is considered low. The option to leave the GBS in place as it is will result in the highest number of species maintained in the local area.

## 6. ACKNOWLEDGEMENTS

We thank the crew of the Cdt. Fourcault (IMO 7304675; NV Seatec). Without their professional support the dives would have been impossible. We are grateful to the supporting divers, including Joost Bergsma (Bureau Waardenburg), Ben Stiefelhagen and Melchior Stiefelhagen (Get Wet Maritiem) and Klaudie Bartelink and Peter van Rodijnen (Dutch Maritime Productions). Emiel Brummelhuis, Jack Perdon and Jan Tjalling van der Wal (Wageningen Marine Research) were involved in our lab work or data-processing and we thank them for their help. The work reported in this publication was funded by Petrogas E&P Netherlands B.V.. We thank Alan Shand and Gert de Raadt at Petrogas for making this work possible. We thank Ruud Schulte and Michiel Harings at EBN for their comments on our analysis and outcome. We thank Prof. em. Han Lindeboom for commenting on our work and providing interesting viewpoints on the topic. An earlier version of this text was reviewed by Oscar Bos (Wageningen Marine Research), we thank him for his help improving the manuscript. We thank our anonymous reviewers for their constructive comments on the earlier versions of this text.

## REFERENCES

Adams, T.P., Miller, R.G., Aleynik, D., Burrows, M.T., 2014. Offshore marine renewable energy devices as stepping stones across biogeographical boundaries. J. Appl. Ecol. 51, 330–338. https://doi.org/10.1111/1365-2664.12207

Anderson, M.J., 2005. PERMANOVA: a FORTRAN computer program for permutational multivariate analysis of variance. Department of Statistics, University of Auckland, Auckland, New Zealand. https://doi.org/10.1139/cjfas-58-3-626

Anderson, M.J., 2001. A new method for non-parametric multivariate analysis of variance. Austral Ecol. 26, 32–46.

Bolam, S.G., Garcia, C., Eggleton, J., Kenny, A.J., Buhl-Mortensen, L., Gonzalez-Mirelis, G., Van Kooten, T., Dinesen, G., Hansen, J., Hiddink, J.G., Sciberras, M., Smith, C., Papadopoulou, N., Gumus, A., Van Hoey, G., Eigaard, O.R., Bastardie, F., Rijnsdorp, A.D., 2017. Differences in biological traits composition of benthic assemblages between unimpacted habitats. https://doi.org/10.1016/j.marenvres.2017.01.004

Bolam, S.G., Mcilwaine, P.S.O., Garcia, C., 2016. Application of biological traits to further our understanding of the impacts of dredged material disposal on benthic assemblages. https://doi.org/10.1016/j.marpolbul.2016.02.031

Bos, O.G., Gittenberger, A., Boois, I.J. de, Asch, M. van, Wal, J.T. van der, J. Cremer, B., Hoorn, van der, Pieterse, S., Bakker, P.A.J., 2016. Soortenlijst Nederlandse Noordzee. Wageningen University & Research Rapport C125/16A. Den Helder.

Boström, C., Törnroos, A., Bonsdorff, E., 2010. Invertebrate dispersal and habitat heterogeneity: Expression of biological traits in a seagrass landscape. J. Exp. Mar. Bio. Ecol. 390, 106–117. https://doi.org/10.1016/j.jembe.2010.05.008

Bray, J.R., Curtis, J.T., 1957. An ordination of the upland forest communities of southern Wisconsin. Ecol. Monogr. 27, 325–349.

Bull, A.S., Love, M.S., 2019. Worldwide oil and gas platform decommissioning: A review of practices and reefing options. Ocean Coast. Manag. 168, 274–306. https://doi.org/10.1016/J.OCECOAMAN.2018.10.024

Calow, P., 1972. A method for determining the surface areas of stones to enable quantitative density estimates of littoral stonedwelling organisms to be made. Hydrobiologia 40, 37–50. https://doi.org/10.1007/BF00123590

Causon, P.D., Gill, A.B., 2018. Linking ecosystem services with epibenthic biodiversity change following installation of offshore wind farms. Environ. Sci. Policy 89, 340–347. https://doi.org/10.1016/j.envsci.2018.08.013

Chao, A., 1987. Estimating the population size for capture-recapture data with unequal catchability. Biometrics 43, 783–791.

Chao, A., Gotelli, N.J., Hsieh, T.C., Sander, E.L., Ma, K.H., Colwell, R.K., Ellison, A.M., 2014. Rarefaction and extrapolation with Hill numbers: a framework for sampling and estimation in species diversity studies. Ecol. Monogr. 84, 45–67. https://doi.org/10.1890/13-0133.1

Coates, D. a., Deschutter, Y., Vincx, M., Vanaverbeke, J., 2014. Enrichment and shifts in macrobenthic assemblages in an offshore wind farm area in the Belgian part of the North Sea. Mar. Environ. Res. 95, 1–12. https://doi.org/10.1016/j.marenvres.2013.12.008

Coolen, J.W.P., Boon, A.R., Crooijmans, R.P., Van Pelt, H., Kleissen, F., Gerla, D., Beermann, J., Birchenough, S.N.R., Becking, L.E., Luttikhuizen, P.C., 2020. Marine stepping-stones: Water flow drives Mytilus edulis population connectivity between offshore energy installations. Mol. Ecol. 29, 686–703. https://doi.org/10.1111/mec.15364

Coolen, J.W.P., Bos, O.G., Glorius, S., Lengkeek, W., Cuperus, J., van der Weide, B.E., Agüera, A., 2015a. Reefs, sand and reef-like sand: A comparison of the benthic biodiversity of habitats in the Dutch Borkum Reef Grounds. J. Sea Res. 103, 84–92.

Coolen, J.W.P., Lengkeek, W., Cuperus, J., van der Weide, B.E., 2016a. Data from: Distribution of the invasive Caprella mutica Schurin, 1935 and native Caprella linearis (Linnaeus, 1767) on artificial hard substrates in the North Sea: separation by habitat. Dryad Digit. Repos. https://doi.org/10.5061/dryad.m563r

Coolen, J.W.P., Lengkeek, W., Degraer, S., Kerckhof, F., Kirkwood, R.J., Lindeboom, H.J., 2016b. Distribution of the invasive *Caprella mutica* Schurin, 1935 and native *Caprella linearis* (Linnaeus, 1767) on artificial hard substrates in the North Sea: separation by habitat. Aquat. Invasions 11, 437–449.

Coolen, J.W.P., Lengkeek, W., Lewis, G., Bos, O.G., van Walraven, L., van Dongen, U., 2015b. First record of Caryophyllia smithii in the central southern North Sea: artificial reefs affect range extensions of sessile benthic species. Mar. Biodivers. Rec. 8, 4 pages. https://doi.org/10.1017/S1755267215001165

Coolen, J.W.P., van der Weide, B.E., Cuperus, J., Blomberg, M., van Moorsel, G.W.N.M., Faasse, M.A., Bos, O.G., Degraer, S., Lindeboom, H.J., 2018. Benthic biodiversity on old platforms, young wind farms and rocky reefs. ICES J. Mar. Sci. fsy092.

Cooper, C.M., Testa, S., 2001. A quick method of determining rock surface area for quantification of the invertebrate community. Hydrobiologia 452, 203–208. https://doi.org/10.1023/A:1011914624264

Dannheim, J., Bergström, L., Birchenough, S.N.R., Brzana, R., Boon, A.R., Coolen, J.W.P., Dauvin, J.-C., De Mesel, I., Derweduwen, J., Gill, A.B., Hutchison, Z.L., Jackson, A.C., Janas, U., Martin, G., Raoux, A., Reubens, J., Rostin, L., Vanaverbeke, J., Wilding, T.A., Wilhelmsson, D., Degraer, S., 2020. Benthic effects of offshore renewables: identification of knowledge gaps and urgently needed research. ICES J. Mar. Sci. 77, 1092–1108. https://doi.org/10.1093/icesjms/fsz018

De Mesel, I., Kerckhof, F., Norro, A., Rumes, B., Degraer, S., 2015. Succession and seasonal dynamics of the epifauna community on offshore wind farm foundations and their role as stepping stones for non-indigenous species. Hydrobiologia 756, 37–50. https://doi.org/10.1007/s10750-014-2157-1

de Vrees, L., 2019. Adaptive marine spatial planning in the Netherlands sector of the North Sea. Mar. Policy 103418. https://doi.org/10.1016/j.marpol.2019.01.007

Dias, I.M., Spierings, M., Coolen, J.W.P., van der Weide, B.E., Cuperus, J., 2017. First record of Syllis vittata (Polychaeta: Syllidae) in the Dutch North Sea. Mar. Biodivers. Rec. 10, 16. https://doi.org/10.1186/s41200-017-0120-3

Ellis, J., Ellis, J., Solander, D.C., 1786. The natural history of many curious and uncommon zoophytes : collected from various parts of the globe. Printed for Benjamin White and Son … and Peter Elmsly …, London :

EMODnet, 2020. European Marine Observation Data Network (EMODnet) Geology portal [WWW Document]. URL https://data.geus.dk/egdi/?mapname=egdi_emodnet_geology&showCustomLayers=true#baslay=baseMapGEUS&optlay=&extent=3273100,2808560,3812610,3078310&layers=emodnet_substrate_multiscale (accessed 1.6.20).

EMODnet, 2019. European Marine Observation Data Network (EMODnet) Bathymetry portal [WWW Document]. URL https://www.emodnet-bathymetry.eu (accessed 12.5.19).

Euler, L., 1768. Lettres a une princesse d’Allemagne sur divers sujets de physique & de philosophie. A Saint Petersbourg: De l’Imprimerie de l’Academie impériale des sciences, Saint Petersbourg.

Faasse, M., Coolen, J.W.P., Gittenberger, A., Schrieken, N., 2016. Nieuwe mosdiertjes van noordzeewrakken (Bryozoa). Ned. Faun. Meded. 46, 43–48.

Fowler, A.M., Jørgensen, A., Svendsen, J.C., Macreadie, P.I., Jones, D.O., Boon, A.R., Booth, D.J., Brabant, R., Callahan, E., Claisse, J.T., Dahlgren, T.G., Degraer, S., Dokken, Q.R., Gill, A.B., Johns, D.G., Leewis, R.J., Lindeboom, H.J., Linden, O., May, R., Murk, A.J., Ottersen, G., Schroeder, D.M., Shastri, S.M., Teilmann, J., Todd, V., Hoey, G. Van, Vanaverbeke, J., Coolen, J.W.P., 2018. Environmental benefits of leaving offshore infrastructure in the ocean. Front. Ecol. Environ. 16, 571–578. https://doi.org/10.1002/FEE.1827

Friedlander, A.M., Ballesteros, E., Fay, M., Sala, E., 2014. Marine communities on oil platforms in Gabon, West Africa: High biodiversity oases in a low biodiversity environment. PLoS One 9. https://doi.org/10.1371/journal.pone.0103709

Fujii, T., 2015. Temporal variation in environmental conditions and the structure of fish assemblages around an offshore oil platform in the North Sea. Mar. Environ. Res. 108, 69–82. https://doi.org/10.1016/j.marenvres.2015.03.013

Glasby, T.M., Connell, S.D., 2001. Orientation and position of substrata have large effects on epibiotic assemblages. Mar. Ecol. Prog. Ser. 214, 127–135. https://doi.org/10.3354/meps214127

Goddard, H.R., Love, M.S., 2010. Megabenthic invertebrates on shell mounds associated with oil and gas platforms off california. Bull. Mar. Sci. 86, 533–554.

Graham, A.A., McCaughan, D.J., McKee, F.S., 1988. Measurement of surface area of stones. Hydrobiologia 157, 85–87. https://doi.org/10.1007/BF00008813

Henry, L.-A., Mayorga-Adame, C.G., Fox, A.D., Polton, J.A., Ferris, J.S., McLellan, F., McCabe, C., Kutti, T., Roberts, J.M., 2018. Ocean sprawl facilitates dispersal and connectivity of protected species. Sci. Rep. 8, 11346. https://doi.org/10.1038/s41598-018-29575-4

Hijmans, R.J., 2019. raster: Geographic Data Analysis and Modeling. R package version 3.0-7.

Holstein, J., 2018. worms: Retriving Aphia Information from World Register of Marine Species. R package version 0.2.2.

IPIECA, 2010. Alien invasive species and the oil and gas industry. Guidance for prevention and management. IPIECA report. London, UK.

Kingdom of the Netherlands, 2020. Mijnbouwwet - BWBR0014168 [WWW Document]. wetten.overheid.nl. URL https://wetten.overheid.nl/BWBR0014168/2020-03-18 (accessed 4.9.20).

Kröncke, I., Reiss, H., Dippner, J.W., 2013. Effects of cold winters and regime shifts on macrofauna communities in shallow coastal regions. Estuar. Coast. Shelf Sci. 119, 79–90. https://doi.org/10.1016/j.ecss.2012.12.024

Krone, R., Gutow, L., Joschko, T.J., Schröder, A., 2013. Epifauna dynamics at an offshore foundation-implications of future wind power farming in the North Sea. Mar. Environ. Res. 85, 1–12. https://doi.org/10.1016/j.marenvres.2012.12.004

Krone, R., Schröder, A., 2011. Wrecks as artificial lobster habitats in the German Bight. Helgol. Mar. Res. 65, 11–16. https://doi.org/10.1007/s10152-010-0195-2

Langeveld, B., Creuwels, J., Slieker, F., van der Es, H., 2020. Natural History Museum Rotterdam - Specimens. Version 1.6. Natural History Museum Rotterdam. Occurrence dataset [WWW Document]. https://doi.org/10.15468/kwqaay

Larsson, J., 2019. eulerr: Area-Proportional Euler and Venn Diagrams with Ellipses. R package version 6.0.0.

Love, M.S., Schroeder, D.M., Nishimoto, M.M., 2003. The ecological role of oil and gas production platforms and natural outcrops on fishes in southern and central california: A synthesis of information, U.S. Geological Survey.

Luttikhuizen, P., Beermann, J., Crooijmans, R., Jak, R., Coolen, J.W.P., 2019. Low genetic connectivity in a fouling amphipod among man-made structures in the southern North Sea. Mar. Ecol. Prog. Ser. 133–142. https://doi.org/10.3354/meps12929

Macreadie, P.I., Fowler, A.M., Booth, D.J., 2011. Rigs-to-reefs: Will the deep sea benefit from artificial habitat? Front. Ecol. Environ. 9, 455–461. https://doi.org/10.1890/100112

Marine Information and Data Centre, 2019. Geoviewer open data [WWW Document]. URL https://www.informatiehuismarien.nl/uk/opendata/ (accessed 12.1.19).

Maslin, E., 2016. Platforms, naturally! [WWW Document]. Offshore Eng. URL https://www.oedigital.com/news/447812-platforms-naturally (accessed 1.31.20).

Mason, N.W.H., Mouillot, D., Lee, W.G., Wilson, J.B., 2005. Functional richness, functional evenness and functional divergence: the primary components of functional diversity. Oikos 111, 112–118.

Mavraki, N., Degraer, S., Moens, T., Vanaverbeke, J., 2020a. Functional differences in trophic structure of offshore wind farm communities: A stable isotope study. Mar. Environ. Res. 157, 104868. https://doi.org/10.1016/j.marenvres.2019.104868

Mavraki, N., Degraer, S., Vanaverbeke, J., Braeckman, U., 2020b. Organic matter assimilation by hard substrate fauna in an offshore wind farm area: a pulse-chase study. ICES J. Mar. Sci. in press.

Naturalis, EIS, 2020. Nederlands Soortenregister [WWW Document]. URL https://www.nederlandsesoorten.nl/ (accessed 4.8.20).

Oksanen, J., Blanchet, F.G., Friendly, M., Kindt, R., Legendre, P., McGlinn, D., Minchin, P.R., O’Hara, R.B., Simpson, G.L., Solymos, P., Henry, M., Stevens, H., Szoecs, E., Wagner, H., 2019. vegan: Community Ecology Package. R package version 2.5-6.

Oshurkov, V. V., 1992. Succession and climax in some fouling communities. Biofouling 6, 1–12. https://doi.org/10.1080/08927019209386205

OSPAR Commission, 1998. OSPAR 98/3 on the Disposal of Disused Offshore Installations.

Picken, G.B., 1986. Moray Firth marine fouling communities. Proc. R. Soc. Edinburgh. Sect. B. Biol. Sci. 91, 213–220.

Picken, G.B., Baine, M., Heaps, L., Side, J., 2000. Rigs to reefs in the North Sea, in: Jensen, A.C., Collins, K.J., Lockwood, A.P.M. (Eds.), Artificial Reefs in European Seas. Kluwer Academic Publishers, Dordrecht, pp. 331–342.

Pradella, N., Fowler, a. M., Booth, D.J., Macreadie, P.I., 2014. Fish assemblages associated with oil industry structures on the continental shelf of north-western Australia. J. Fish Biol. 84, 247–255. https://doi.org/10.1111/jfb.12274

R Core Team, 2019. R: A language and environment for statistical computing (Version 3.6.1).

Reubens, J.T., Degraer, S., Vincx, M., 2011. Aggregation and feeding behaviour of pouting (Trisopterus luscus) at wind turbines in the Belgian part of the North Sea. Fish. Res. 108, 223–227. https://doi.org/DOI10.1016/j.fishres.2010.11.025

Reubens, J.T., Vandendriessche, S., Zenner, A.N., Degraer, S., Vincx, M., 2013. Offshore wind farms as productive sites or ecological traps for gadoid fishes? - Impact on growth, condition index and diet composition. Mar. Environ. Res. 90, 66–74. https://doi.org/10.1016/j.marenvres.2013.05.013

RStudio, 2019. RStudio: Integrated development environment for R (Version 1.2.5001).

Russell, D.J.F., Brasseur, S.M.J.M., Thompson, D., Hastie, G.D., Janik, V.M., Aarts, G., McClintock, B.T., Matthiopoulos, J., Moss, S.E.W., B. McConnell, 2014. Marine mammals trace anthropogenic structures at sea. Curr. Biol. 24, 638–639. https://doi.org/10.1016/j.cub.2014.06.033

Ryland, J.S., Holt, R., Loxton, J., Jones, M.E.S., Porter, J.S., 2014. First occurrence of the non-native bryozoan Schizoporella japonica Ortmann (1890) in Western Europe. Zootaxa 3780, 481–502. https://doi.org/10.11646/zootaxa.3780.3.3

Schroeder, D.M., Love, M.S., 2004. Ecological and political issues surrounding decommissioning of offshore oil facilities in the Southern California Bight. Ocean Coast. Manag. 47, 21–48. https://doi.org/10.1016/j.ocecoaman.2004.03.002

Simpson, E.H., 1949. Measurement of diversity. Nature. https://doi.org/10.1038/163688a0

van der Molen, J., García-García, L.M., Whomersley, P., Callaway, A., Posen, P.E., Hyder, K., 2018. Connectivity of larval stages of sedentary marine communities between hard substrates and offshore structures in the North Sea. Sci. Rep. 8, 14772. https://doi.org/10.1038/s41598-018-32912-2

van der Stap, T., Coolen, J.W.P., Lindeboom, H.J., 2016. Marine fouling assemblages on offshore gas platforms in the southern North Sea: Effects of depth and distance from shore on biodiversity. PLoS One 11, 1–16. https://doi.org/10.1371/journal.pone.0146324

Vance, T., Lauterbach, L., Lenz, M., Wahl, M., Sanderson, R. a., Thomason, J.C., 2009. Rapid invasion and ecological interactions of Diplosoma listerianum in the North Sea, UK. Mar. Biodivers. Rec. 2, 1–5. https://doi.org/10.1017/S1755267209000815

Whomersley, P., Picken, G.B., 2003. Long-term dynamics of fouling communities found on offshore installations in the North Sea. J. Mar. Biol. Assoc. U.K. 83, 897–901. https://doi.org/10.1017/S0025315403008014h

Wilhelmsson, D., Malm, T., 2008. Fouling assemblages on offshore wind power plants and adjacent substrata. Estuar. Coast. Shelf Sci. 79, 459–466. https://doi.org/DOI10.1016/j.ecss.2008.04.020

WoRMS Editorial Board, 2019. World Register of Marine Species [WWW Document]. URL http://www.marinespecies.org (accessed 12.1.19).

Yeo, D.C., Ahyong, S.T., Lodge, D.M., Ng, P.K., Naruse, T., Lane, D.J., 2010. Semisubmersible oil platforms: understudied and potentially major vectors of biofouling-mediated invasions. Biofouling 26, 179–186. https://doi.org/10.1080/08927010903402438

Zintzen, V., 2007. Biodiversity of shipwrecks from the Southern Bight of the North Sea. University of Louvain, Belgium.

Zintzen, V., Massin, C., 2010. Artificial hard substrata from the Belgian part of the North Sea and their influence on the distributional range of species. Belgian J. Zool. 140, 20–29.

Zintzen, V., Norro, A., Massin, C., Mallefet, J., 2007. Temporal variation of Tubularia indivisa (Cnidaria, Tubulariidae) and associated epizoites on artificial habitat communities in the North Sea. Mar. Biol. 153, 405–420. https://doi.org/10.1007/s00227-007-0819-5

